# Updated Metabolome Annotation Reveals Plasma and Fecal Metabolic Signatures Modulated by Microbiota Transplant Therapy in Autism Spectrum Disorder

**DOI:** 10.1101/2024.12.16.628826

**Authors:** Khemlal Nirmalkar, Fatir Qureshi, Dae-Wook Kang, Manish Muchapothula, Evelyn Takyi, Blake Dirks, Christina K. Flynn, Juergen Hahn, James B. Adams, Rosa Krajmalnik-Brown

## Abstract

While Autism Spectrum Disorder (ASD) is diagnosed through behavioral symptoms and psychometric evaluations, it has also been associated with distinct metabolomic patterns. A previous clinical trial of Microbiota Transplant Therapy (MTT) in children with ASD and gastrointestinal (GI) issues revealed significant differences in plasma metabolomics between children with ASD and their typically developing (TD) counterparts, which diminished after MTT. The objective of this study was to reanalyze the plasma and fecal samples using updated metabolomics libraries at Metabolon together with a comprehensive panel of statistical methods to provide deeper insights into ASD-related metabolic differences and the impact of MTT. Compared with the original analysis, the updated annotation identified substantially more annotated metabolites and uncovered additional plasma and fecal metabolic changes associated with MTT. The reanalysis highlighted specific metabolites whose relative peak intensities differed between the ASD and TD groups, as well as metabolites with significant changes following MTT. Several plasma metabolites, including sarcosine, iminodiacetate, caproate, and caprylate, initially showed significant differences between the ASD and TD groups but shifted to resemble TD levels after MTT. In fecal samples, *p*-cresol sulfate and sphingolipids emerged as metabolites with altered intensities following MTT. Multivariate Fisher’s Discriminant Analysis (FDA) with leave-one-out cross-validation revealed that a set of metabolites including *p*-cresol sulfate, hydroxyproline, and caprylate could robustly classify the ASD and TD cohorts before treatment. However, after treatment, the same FDA model could no longer distinguish the two groups, as FDA scores became similar to those of the TD cohort.

These findings improve our understanding of ASD-associated metabolic alterations and demonstrate that reanalysis of existing untargeted metabolomics datasets using updated metabolite annotation resources and complementary statistical approaches can reveal biologically relevant metabolite changes that were not detected previously. Larger studies with placebo-controlled designs are needed to validate these findings, further define the underlying biochemical pathways, and evaluate their potential for developing personalized therapeutic strategies.

**IMPORTANCE:** Reanalysis of existing untargeted metabolomics datasets using updated metabolite annotation resources and complementary statistical approaches can reveal biological insights that were not apparent in earlier analyses. By leveraging updated metabolomics libraries and expanded statistical analyses, we identified additional annotated metabolites and uncovered plasma and fecal metabolic changes associated with autism spectrum disorder (ASD) and Microbiota Transplant Therapy (MTT). Several metabolites, including sarcosine, caprylate, and *p*-cresol sulfate, distinguished children with ASD from typically developing (TD) controls at baseline and shifted toward TD-like profiles following MTT. These findings demonstrate the value of periodically reanalyzing legacy metabolomics datasets as annotation resources improve, providing a more comprehensive understanding of disease-associated metabolic alterations and therapeutic responses.

## Introduction

In the past decade, the prevalence of autism spectrum disorder (ASD) in the United States has significantly increased, with 1 in 31 children now diagnosed (CDC, 2025). Individuals with ASD experience social communication and behavioral impairments (Adams et al., 2011; Lord et al., 2018), and often have persistent gastrointestinal (GI) issues, such as constipation and diarrhea (Adams et al., 2011; Hung et al., 2023). These co-occurring conditions correlate with increased ASD behavioral severity (Adams et al., 2011), prompting research intersecting their relationship with the gut-brain axis and gut microbiome. Studies reveal significant differences in gut microbiome compositions between ASD individuals and typically developing (TD) children, including higher levels of *Clostridia* species and lower levels of beneficial microbes like *Bifidobacterium* (Kang et al., 2019; Morton et al., 2023).

The complex interaction among microbes, influenced by their metabolites, has been hypothesized to affect GI and behavioral symptoms in ASD (Adams et al., 2011; Krajmalnik-Brown et al., 2014; Kang et al., 2020; Needham et al., 2021; Morton et al., 2023; Hung et al., 2023; Zheng et al., 2023). Differences in metabolite profiles in fecal, urine, and blood samples have been reported in ASD individuals compared to TD peers (Gabriele et al., 2014; Kang et al., 2018; 2020; Needham et al., 2021; Zheng et al., 2021). For example, elevated levels of fecal short-chain fatty acids and blood glutamate, along with increased blood gamma-aminobutyric acid (GABA), have been observed in people with ASD (Shinohe et al., 2006; Kang et al., 2018; El-Ansary et al., 2014; Khalifa et al., 2019). GABA and glutamate, which are critical for cognition and mood regulation, show an altered levels in ASD individuals, suggesting an underlying neuroinflammation (El-Ansary et al., 2014). Additionally, other gut-derived metabolites, such as p-cresol and p-cresol sulfate, are found at higher levels in ASD individuals’ urine, feces, or serum (Flynn et al., 2025; Kang et al., 2018; De Angelis et al., 2013; Gabriele et al., 2014).

Given the microbiome-gut-ASD interconnection, Microbiota Transplant Therapy (MTT) has been explored for its potential benefits in ASD (Kang et al., 2017). Microbiota Transplant Therapy, (redefined from microbiota transfer therapy, Takyi et al. 2025) has shown promise in addressing GI issues and reducing behavioral severity in ASD children. An open-label study by Kang et al. reported an 80% reduction in GI symptoms and a 24% decrease in core ASD symptoms post-MTT, with sustained improvements over two years (Kang et al., 2017, 2019). Increases in beneficial microbes such as *Bifidobacterium*, *Prevotella*, and *Desulfovibrio* were also noted (Kang et al., 2017, 2019).

Building on these findings, untargeted metabolomics of fecal and plasma samples from the same individuals in the MTT study identified several plasma metabolites significantly lower in the ASD group at baseline, including nicotinamide riboside, IMP, iminodiacetate, methylsuccinate, galactonate, valylglycine, sarcosine, and leucylglycine, while caprylate and heptanoate were higher (Kang et al., 2020). Though no fecal metabolite showed significant baseline differences, *p*-cresol sulfate levels were non-significantly higher in ASD children and decreased after MTT (Kang et al., 2020). Multivariate analysis highlighted significant metabolite changes post-MTT, with improvements in sarcosine and inosine 5’-monophosphate in plasma (Adams et al., 2019) and a model of indole, imidazole, propionate, adenosine, and hydroxyproline achieving 96% accuracy in distinguishing ASD from TD children (Qureshi et al., 2020).

In this study, we reanalyzed the metabolomics data collected for the Kang et al. 2017 and 2019 studies. We build upon the findings reported by Kang et al. (2020) by expanding the scope of metabolomics data analyzed through Metabolon’s updated metabolomics libraries and robust statistical methods. Utilizing more up-to-date algorithms and libraries for metabolite detection, we offer a comprehensive comparison of metabolite profiles, encompassing nucleotides, neurotransmitters, amino acids, xenobiotics, and lipids derived from human plasma and fecal samples collected from a carefully matched cohort of individuals with ASD and typically developing children. Previously, data was analyzed using univariate tests and principal-component analysis (PCA) and correlation analysis for selective metabolites. In this reanalysis, we not only replicated the prior analysis techniques, but we also explored other methods such as contrasting the area under the receiver operating characteristic curve (AUROC) between groups, random forest, and Fisher discriminant analysis (FDA) to revalidate the findings (Fig. 1). The results reveal significant differences in metabolite levels between TD and ASD baseline groups in plasma and feces, which were previously unidentified, and important changes shifting the ASD metabolome towards TD after MTT. Together, these findings demonstrate that advances in metabolite annotation and statistical analysis can extract additional biological information from existing untargeted metabolomics datasets, revealing metabolite changes that were not detected in the original analysis.

**Figure 1:**
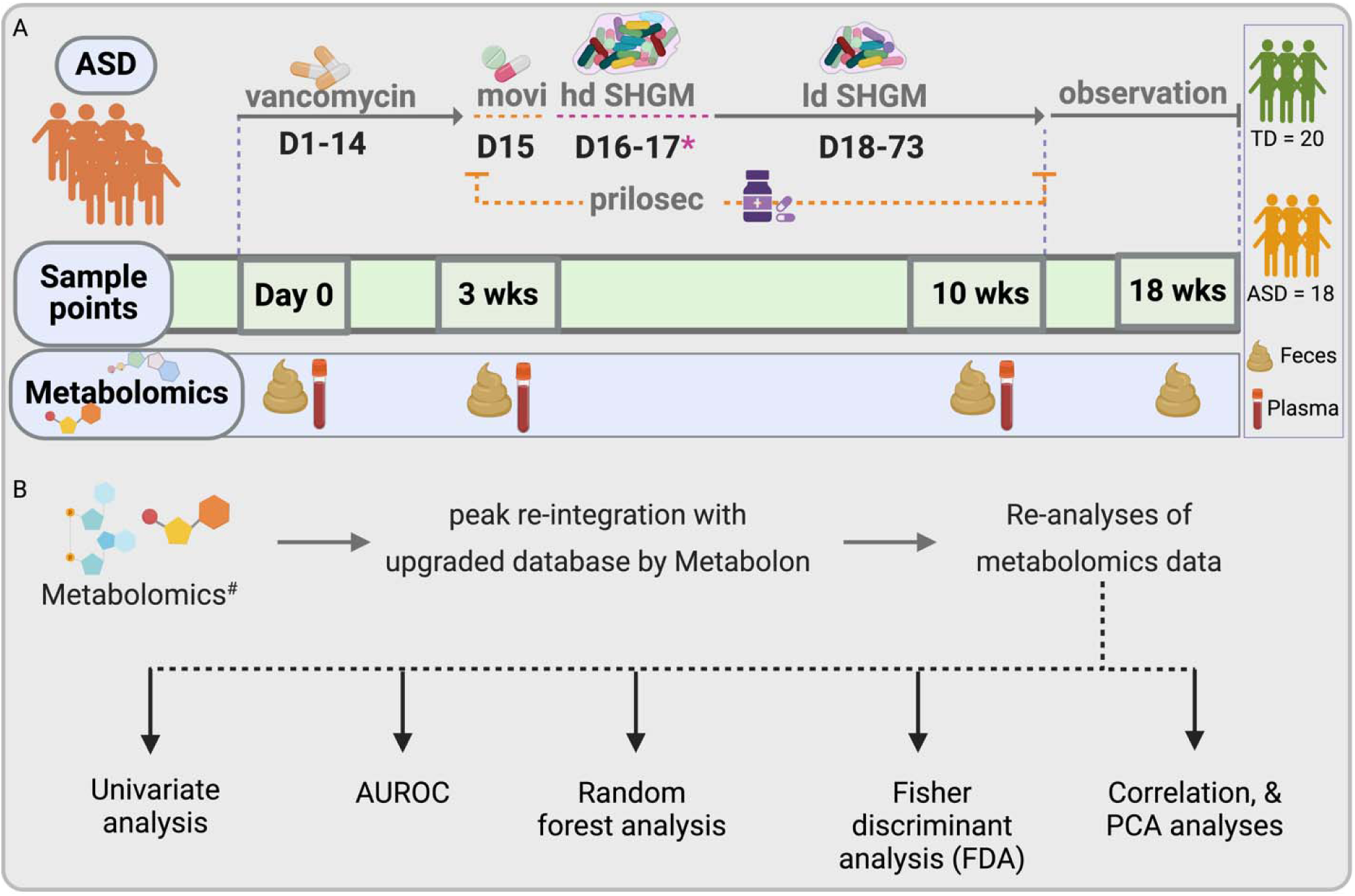
The study design of microbiota transplant therapy for fecal and plasma metabolomics (A) and the overview of the data analyses for fecal and plasma metabolites (B). Plasma was collected right before high-dose microbiota and feces were collected right after high-dose microbiota.

## Methods

### Study design, sample collection and data processing

This study extends the work of Kang et al. (2020) by performing a more thorough reanalysis of plasma and fecal metabolomics data collected before and after MTT in children with ASD. The original protocol was approved by the U.S. Food and Drug Administration (FDA) (investigational new drug number (IND) 15886) and the Institutional Review Board of Arizona State University (ASU IRB protocols #00001053 and #00004890). The trial was registered at ClinicalTrials.gov (NCT02504554), as described in Kang et al. (2017, 2020). Study participants, consent, MTT, and sample collections were described in detail in our previous studies (Kang et al., 2017; 2020). In brief, 20 typically developing (TD) children and eighteen children with ASD were recruited, in the age range of 7-16 years old. Children with ASD participated in an open-label clinical trial including vancomycin for 2 weeks, one-day bowel cleanse with MoviPrep, followed by 7-8 weeks of MTT and Prilosec (a stomach acid suppressant), and a follow-up at 8 weeks post-treatment (i.e., 18 weeks from Day 0) (Kang et al., 2017).

For metabolomic analysis, we collected plasma samples at ASD baseline, week3 (after the vancomycin and bowel cleanse and just before the first dose of microbiota), and week10 (the end of MTT treatment) (Fig. 1). We had one missing plasma sample in week3. Fecal samples were collected at baseline, week3 (right after an initial dose of microbiota), week10 and week18 (8 weeks after the end of MTT). For TD children, we collected plasma and feces only at baseline as they were not treated. All samples were frozen immediately after collection, shipped in dry ice to ASU, and stored at -80°C. Demographic, medical data and diet history was recorded, as described by Kang *et al*., 2017, 2020. Aliquoted frozen samples were analyzed by Metabolon Inc. (Durham, NC, USA) using ultrahigh-performance liquid chromatography-tandem mass spectroscopy (UHPLC-MS/MS) instruments for untargeted metabolomics.

This reanalysis leverages upgrades in peak identification algorithms, software, and libraries available to Metabolon Inc. (https://www.metabolon.com/). In feces, a total of 1,068 metabolites/sample were detected across all samples. Of these, 825 metabolites (77.3%) were chemically identified and assigned a metabolite name, while the remaining 243 features (22.7%) were unnamed. Similarly, in the plasma samples, a total of 1,001 metabolites were detected, with 844 (84.3%) chemically identified and 157 (15.7%) remaining unnamed.

Their proprietary software integrates the mass spectrometry data with retention time alignment, isotope pattern analysis, and database matching to enhance metabolite annotation accuracy. In total, 5,400 compounds were detectable via untargeted metabolomics, with available annotations. These compounds in turn span more than 70 major biological pathways and cover the main classes of endogenous as well as exogenous compounds. Raw data (Supplemental material SM1) underwent imputation and z-score normalization according to the methods described by Kang et al., 2020. Using these extended approaches new data were generated, resulting in modified abundances of metabolites (peak area/intensity) and the identification of additional metabolites within the samples. To focus the analysis on identified compounds for biological interpretation, metabolites lacking chemical annotations (e.g., unidentified compounds labeled as ‘X-###’) were excluded from downstream analysis.

It is important to note that, we recorded xenobiotics ∼16% in plasma and ∼18% in feces which was 1% higher than our previous study (Kang et al., 2020). The slight increase likely reflects updates in Metabolon’s annotation libraries during reanalysis. Importantly, Metabolon’s xenobiotic category encompasses a broad range of compounds, including not only exogenous chemicals (e.g., drugs, pollutants) but also microbial co-metabolites, dietary constituents, and plant-derived compounds. For instance, gut-derived metabolites such as p-cresol and 4-ethylphenyl sulfate are classified by metabolon as xenobiotics due to their chemical features and involvement in pathways like benzoate metabolism. Therefore, the relatively high proportion of xenobiotics observed here may reflect the microbiome-influenced metabolic landscape of individuals with autism, rather than an overrepresentation of exogenous exposures.

### Univariate analysis

Initially, univariate hypothesis testing was employed to compare metabolites between groups. Depending on the data distribution and type of relationship of the two groups, the most appropriate statistical test was selected. Mann-Whitney was utilized for comparing non-parametric data with similar distributions between the TD and ASD samples, while a t-test was used in alternative cases. When comparing ASD groups at various time points a Wilcoxon sign-rank test was used for non-parametric distributions and a paired t-test was used for parametric distributions. False discovery rates (FDR) were determined using the leave-one-out (LOO) approach. This technique is well-suited for high-dimensional data that has an underlying correlational structure, as is common in metabolic studies (Arici et al., 2025). This method applies the same hypothesis testing approach to every combination of a dataset with one sample excluded. Then, the proportion of significant findings is determined to evaluate the likelihood of false discovery. Subsequently, findings with p < 0.05 and FDR <0.05 were considered statistically significant.

### Area Under the Receiver Operating Characteristic Curve (AUROC)

The area under the receiver operating characteristic curve (AUROC) was used to assess the degree to which metabolite measurements were discernible between pre-treatment and post-treatment time points. For a binary classification task, the AUROC is determined by plotting the true positive rate (TPR) against the false positive rate (FPR) of a model across varying threshold values. The area under the TPR/FNR plot corresponds to the AUROC and can serve as a metric to quantify the degree to which different groups are separable. The AUROC value ranges from 0 to 1, where a value of 0.5 indicates no discrimination, a value of 1 represents perfect discrimination, and a value close to 0 suggests that the model is performing worse than random, potentially predicting the opposite of the true outcome.

The AUROC was calculated by treating the measurements of each metabolite individually as the classifier input, which was subject to a threshold value. AUROC was determined for each fecal and plasma metabolite, with baseline measurements being compared against all other time points. The measurements collected from the ASD group were also compared against the TD measurements, both in their entirety and on a time-point basis. By assessing all the AUROC values generated for each metabolite between baseline, week3, and week10 vs TD across, the average fecal and plasma AUROC values for each group were determined.

### Fisher’s Discriminant Analysis (FDA)

FDA is a dimensionality reduction technique that aims to find a transformation of the original data that accentuates the differences between distinct groups while reducing the variations within each group. This method projects multidimensional data into a lower dimensional space in such a way as that dissimilar groups are maximally separated, while members of the same group remain closely grouped together. By doing so, FDA facilitates the creation of more discriminative representations (Huberty, 1975).

Metabolites that had been deemed to be statistically significant in both plasma and feces were used to develop an FDA model for the purpose of classification between ASD and TD cohorts. To mitigate the risk of overfitting, feature selection was limited to five variables, as literature suggests this approach optimally suits small sample sizes with similar data structure properties in ASD prediction. This decision is also supported by the success of prior fecal and plasma model panels achieving the upper bounds of their predictive accuracy with this number of features (Howsmon et al., 2017; Vargason et al., 2019; Qureshi et al., 2020). An exhaustive search of all potential FDA model combinations of five metabolites was determined, using only the ASD baseline and TD groups as training data. For all possible combinations of five metabolites, the AUROC was evaluated to determine whether to conduct further testing. Each metabolite combination model that achieved an AUROC greater than 0.97 was subjected to a leave-one-out cross-validation (LOO-CV) paradigm. The top-performing model, as quantified by CV accuracy, was then used to predict the classification of the MTT week3 and week10 groups.

### Random Forest Analysis

Random forest (RF) is defined as an ensemble learning method that can be used for classification. An RF approach to classification operates by aggregating the collective performance of multiple decision trees and using the plurality of outcomes to predict the group label of a sample. This algorithm is constructed by first setting aside one-third of the training data to utilize as testing data for cross-validation purposes. Subsequently, a user-specified number of decision trees is constructed. The performance of the algorithm is then quantified based on the classification accuracy on the randomly selected test set. A random forest model was fit using all significant metabolomic measurements in order to evaluate the mean decrease in accuracy for the exclusion of each variable. This metric is commonly utilized to provide a perspective on feature importance, which can be used to more efficiently identify variables better suited for distinguishing between labeled categories (Altman et al., 2017). Although random forests are generally more effective with larger datasets, they can still serve as exploratory tools for feature selection in smaller datasets, provided their limitations are acknowledged and results are interpreted cautiously (Wouter et al., 2013)

### Correlation Coefficient

The Pearson‘s correlation coefficient is defined by the strength and direction of the linear relationship between two continuous variables. The Pearson’s correlation coefficient was determined between all metabolite, pathway, and taxa pairs. The significance of the correlation coefficients was evaluated using hypothesis testing with a significance level set at α = 0.05. The robustness of significant findings was cross validated using a LOO approach. Employing this strategy, the significance of the correlation coefficient was determined for all possible combinations of the input measurement with a single sample removed. If a false discovery rate (FDR) of less than 0.05 was determined, the finding was deemed to be significant.

All univariate analysis, area under the receiver operating characteristic (AUROC), random forest analysis and plots were made with Python (v3.8.5) in Jupyter Notebook (v6.1.4) using numpy (v1.19.2), matplotlib (v3.3.2), seaborn (v0.11.0), pandas (v1.1.3) and scipy (v1.5.2). PCA plot was made using MetaboAnalyst v6.0 (https://www.metaboanalyst.ca/). BioRender (Biorender, 2022, Licensed, https://biorender.com) and Inkscape (v1.1) were used to create or edit the figures.

## Results

In the previous study (Kang et al., 2020), 621 plasma and 666 fecal metabolites were identified in the collected samples. Reanalysis of the metabolomics datasets using the updated Metabolon platform increased overall feature detection and metabolite annotation. A total of 1,001 plasma and 1,068 fecal features were detected, of which 84.3% (n=844) in plasma and 77.3% (n=825) in feces were chemically annotated. This represents an increase of 223 plasma and 159 fecal annotated metabolites compared to the original dataset (Kang et al., 2020) (Fig. 2). Unannotated features were excluded from downstream analyses.

**Figure 2.**
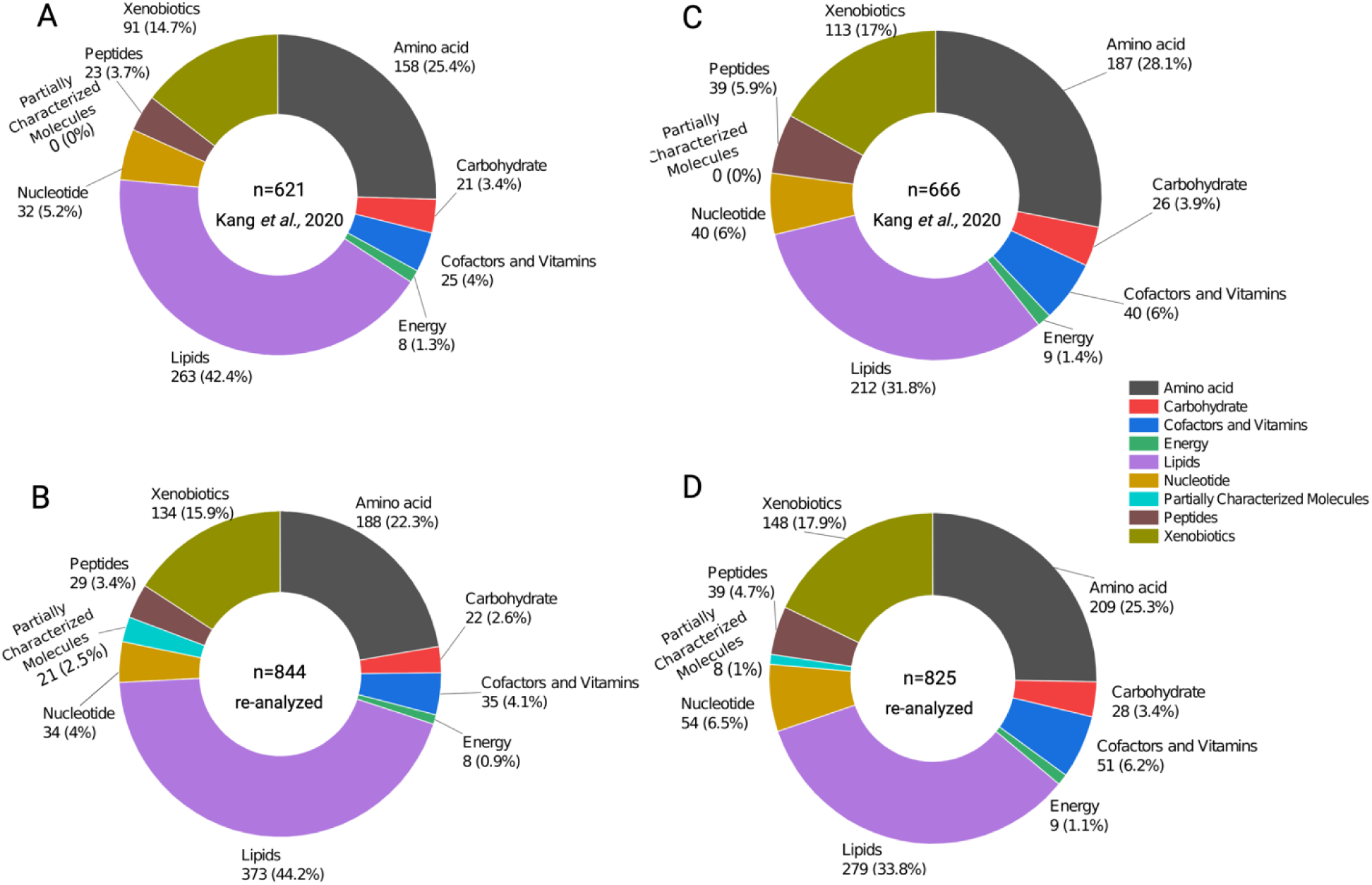
An overview of plasma metabolomics with the previous A) and re-analyzed B), fecal metabolomics with the previous C) and re-analyzed D) of peaks. The pie charts show the total number of metabolites identified and the type of metabolites in plasma and fecal samples.

Metabolon assigned each identified metabolite to a major chemical class (superpathways), including lipids, amino acids, xenobiotics, carbohydrates, nucleotides, cofactors and vitamins, energy metabolites, and peptides. Across both plasma and fecal datasets, the metabolome was dominated by lipids, amino acids, and xenobiotics. Lipids accounted for approximately 44% of plasma metabolites and 34% of fecal metabolites, amino acids for ∼22-25%, and xenobiotics for ∼16-18%. These distributions were broadly consistent between the original and reanalyzed datasets, with modest increases in lipid and xenobiotic representation in the reanalysis (Fig. 2; supplemental material SM1).

To assess the consistency between the original and reanalyzed datasets, we compared the statistical significance of superpathway metabolites within each sample type at each study time point (baseline, 3 weeks, 10 weeks, and 18 weeks, TD). In plasma, a high proportion of metabolites remained significantly different across all time points (∼90-100%), indicating strong agreement between the two analyses. In contrast, fecal metabolomics showed greater variability, with approximately 79% of metabolites remaining significant different at baseline, 7.8% at 3 weeks, 76% at 10 weeks, and 13.9% at 18 weeks. At the global level, principal component analysis and distributional assessments showed separation across sample groups and time points, with clearer structure observed in the reanalyzed dataset (Supplementary file 1 Fig. S1-S2). Detailed results, including superpathway and subpathway summaries, matched metabolite comparisons, and full datasets, are provided in the supplemental materials SM2-SM5.

### Metabolite profiles changed after MTT in ASD children

Univariate analyses showed that certain metabolites that were initially different between ASD and TD groups at baseline, and shifted following MTT. These metabolites relative peak intensity changed post-treatment, and some of these intensities became similar to those in TD in plasma (Fig. 3A, C) and fecal samples (Fig. 3B, Fig. S3). In plasma, out of 844 metabolites, we observed 20 metabolites whose intensity was significantly different between the ASD baseline and TD and that also changed after MTT (Fig. 3A). We grouped these plasma metabolites into two clusters; **1)** Cluster P-high (cluster of 14 plasma metabolites), which includes metabolites that had significantly higher intensity (adjusted *p*<0.05) at ASD baseline vs TD, but after MTT, the relative intensity of these metabolites decreased and became similar to TD (Fig. 3A). Figure 3A shows 12 plasma metabolites in this category, including: glycerophosphoethanolamine (GPEA), caprylate (8:0), and caproate (6:0). The other two plasma metabolites, taurine and adenosine-5’-monophosphate (AMP), were non-significantly higher (*p*>0.05) at baseline vs TD but significantly decreased after MTT (Fig. 3A). **2)** Cluster P-low (cluster of 6 plasma metabolites) which includes metabolites that had significantly lower intensity (adjusted p<0.05) at baseline vs TD, but after MTT, relative intensity of these metabolites significantly increased (adjusted *p*<0.05) and became similar to TD. Cluster P-low includes sarcosine, iminodiacetate, bilirubin, 2-methylserine, and indole propionate.

**Figure 3.**
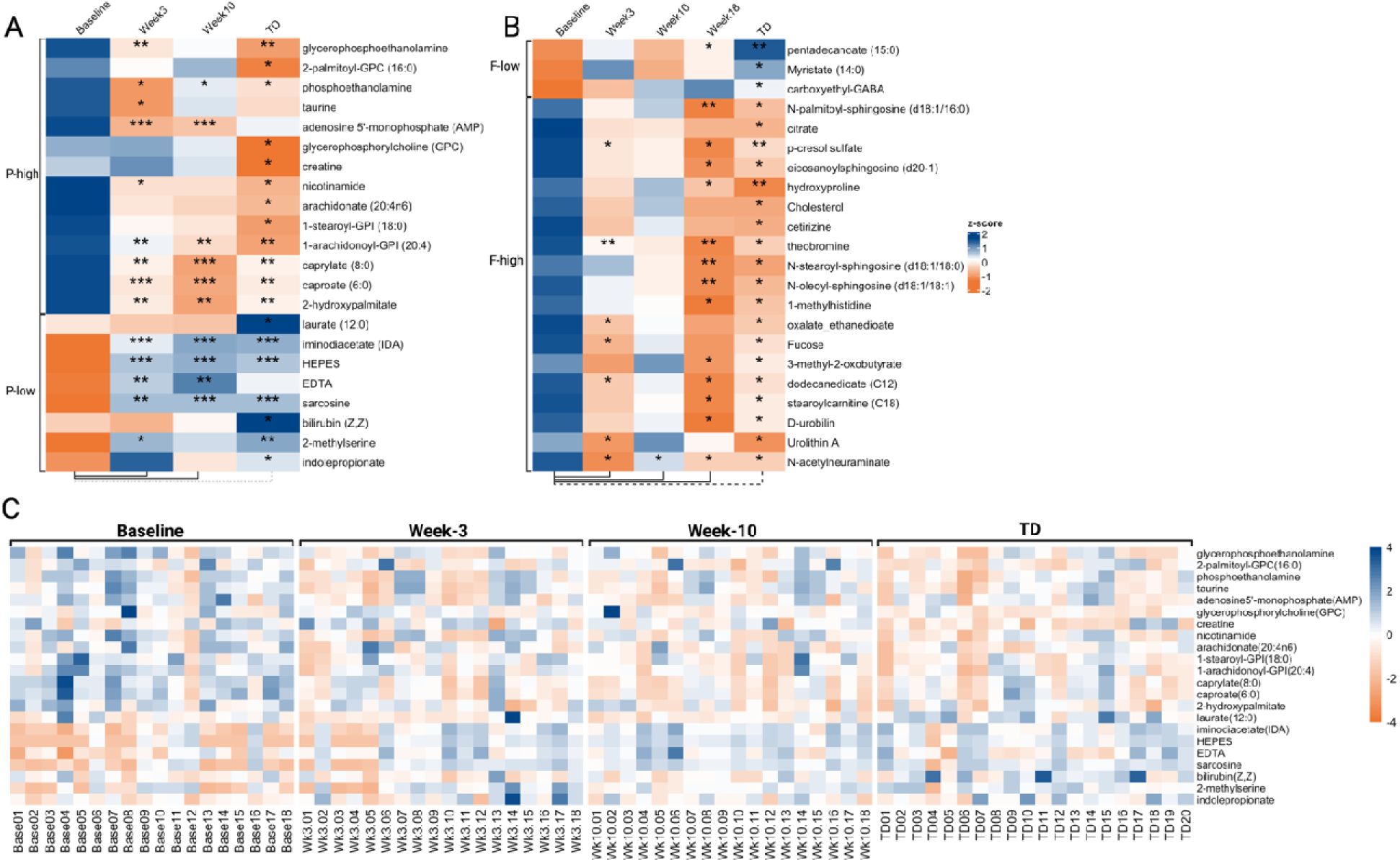
Univariate comparison of the (z-score) relative intensity of plasma (A) and fecal (B) metabolites in children with ASD and comparisons after MTT and with TD. For the plasma metabolite (A), a longitudinal paired-wise univariate analysis was performed at baseline vs. week3, week10 and cross-sectional unpaired comparison vs. TD. Similarly, for the fecal metabolites (B), a longitudinal paired-wise univariate analysis was performed (shown in solid blue lines at the bottom of the heatmap) at baseline vs. week3, week10, week18 and cross-sectional unpaired comparison vs. TD (shown in dashed blue line). (C) heatmap (z-score) for each participant with 20 plasma metabolites that were significantly changed or were different in children with ASD at baseline compared with MTT and with TD groups in Figure 3A. Week18 time was not available for the plasma. See Supplementary Fig. S1 for fecal metabolites heatmap for each participant. The median of relative intensities in plasma (A) and feces (B) was used to construct the heatmap for each group. *Single asterisk indicates p <0.05, **double asterisks indicate p <0.01, triple *** asterisks indicate p <0.001. # indicates the statistically significant (p<0.05) metabolites after log-transformation. All p-values are FDR corrected. ASD: Autism Spectrum Disorders, TD: Typically Developing.

Similar to plasma, out of 825 fecal metabolites, we observed 22 metabolites that had significantly different intensities from TD or changed after MTT (Fig 3B, Fig. S3). Fecal metabolites were also divided into two clusters; **1)** Cluster F-low (cluster of 3 fecal metabolites) where all metabolites had significantly lower intensity (adjusted *p*<0.05) at baseline vs. TD, and **2)** Cluster F-high (cluster of 19 fecal metabolites) where all metabolites had higher intensity (adjusted *p*<0.05) at baseline vs. TD. Cluster F-low (Fig. 3B-) shows 3 fecal metabolites pentadecanoate (15:0), myristate (14:0), and carboxyethyl-GABA whose intensity was significantly lower at baseline vs. TD, and only one metabolite pentadecanoate (15:0) whose intensity significantly increased (adjusted *p*<0.05) after week18 (Fig. 3B). In cluster F-high, 19 metabolites had significantly higher intensity (adjusted *p*<0.05) at baseline vs TD, here the intensity of 5 metabolites: *p*-cresol sulfate, theobromine, oxalate ethanedioate, dodecanedioate (C12), and Urolithin-A significantly decreased (adjusted *p*<0.05) at week3 vs baseline (Fig. 3B). The intensity of N-palmitoyl-sphingosine (d18:1/16:0), *p*-cresol sulfate, eicosanoyl-sphingosine (d20:1), hydroxyproline, theobromine, N-stearoyl-sphingosine (d18:1/18:0), N-oleoyl-sphingosine (d18:1/18:1), 3-methyl-2-oxobutyrate, dodecanedioate (C12), and D-urobilin significantly decreased after week18 compared to baseline. We did not observe any significant change for week10 vs. baseline (Fig. 3B).

### Analytics and Patterns of Differential Metabolomic Composition Appear Similar between Feces and Plasma

We utilized AUROC, PCA analysis and random forest analysis to categorize the differences in metabolomic composition across groups and timepoints. AUROC was calculated for all 20 plasma and 22 fecal metabolites that were significantly different between ASD baseline and TD. To examine how metabolites differed post-MTT, AUROC was recalculated at all time points between the ASD and TD cohorts. The average AUROC for all 20 plasma metabolites was 0.73 comparing ASD baseline vs TD, and ASD baseline vs week3, and week10 AUROC’s were 0.70 and 0.62 respectively (Fig. 4A). When we compared the MTT group with TD, after week3 and week10, we observed that the plasma profile became more similar to TD with AUROC 0.56 for week3 vs TD and 0.59 for week10 vs TD (Fig. 4A). A similar trend was also observed for 22 fecal metabolites, which were initially observed with an average AUROC of 0.74 for ASD baseline vs TD. After MTT, fecal metabolite profiles became more similar to the TD group, with AUROC values of 0.58 for week3 vs TD and 0.61 for week10 vs TD (Fig. 4B). Overall, the observed shift in AUROC values for plasma and fecal metabolites indicates that after MTT, the metabolic composition in plasma and fecal samples from children with ASD became more similar to their TD counterparts.

**Figure 4.**
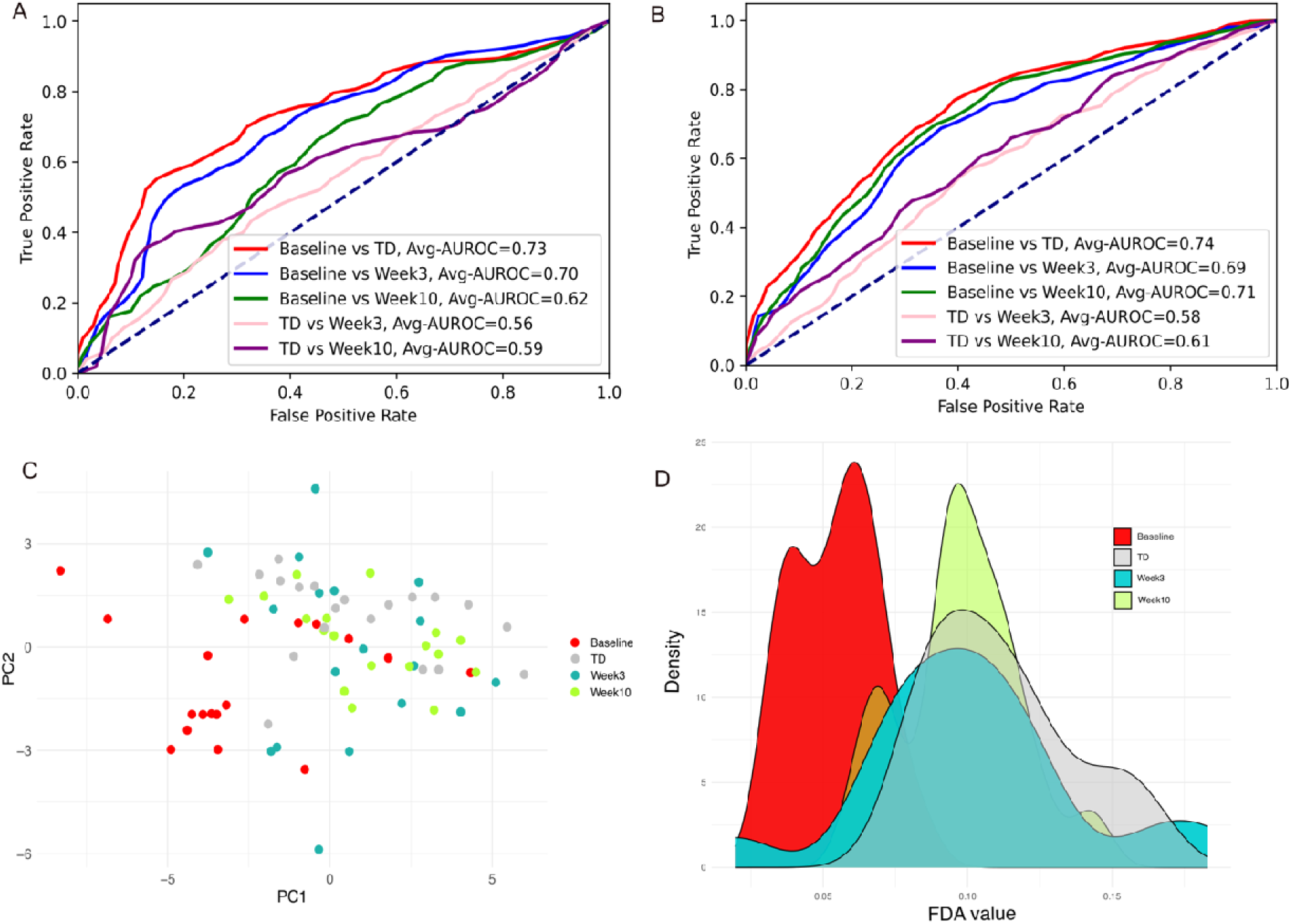
An average Receiver operating characteristic curves (AUROC) of A) plasma, B) fecal metabolites, C) principal component analysis (PCA) of plasma metabolites, and D) FDA analysis of fecal and plasma metabolites (*p*-cresol sulfate, caprylate (C:8) bilirubin, indolepropionate and hydroxyproline) using probability density curves associated with FDA scores before and after MTT in ASD, and TD children. For FDA analysis, true positive vs false positive rate was determined across all threshold values to characterize between ASD Baseline against all possible alternative groups. For AUROC and PCA plots, 22 fecal and 20 plasma metabolites (presented in Figure 3) were used. Refer to Supplementary Figure S2 for PCA plot for fecal metabolites. ASD – autism spectrum disorder, MTT - microbiota transplant therapy, TD - typically developing, FDA - Fisher discriminant analysis.

PCA analysis showed that relative intensity of the 20 significant plasma metabolites at baseline were distinct from TD, but after MTT, became much more similar to TD (Fig. 4C). Fecal samples exhibited greater variance in the typically developing group compared to their ASD counterparts in the first two principal components, and this pattern was also observed in the post treatment ASD group (Fig. S4). Random forest was used as a complimentary method to validate the assessment of each metabolites relative utility in distinguishing between cohorts. Using the full list of metabolites, it was possible to achieve accuracies greater than 0.95. The mean decrease in accuracy for the ASD status prediction model indicated that both fecal and plasma metabolites had a similar influence on model performance (Fig. S5). The top 10 features contributing to mean decrease in accuracy were split evenly between the plasma and fecal metabolites, potentially offering complementary insights into discerning ASD and TD samples. (Fig. S6).

### MTT changed metabolites abundance in ASD towards TD

We selected four plasma metabolites that were highly significantly different at baseline and showed changes after MTT (Fig. 5). A lysine derivative sarcosine and iminodiacetate (IDA) were significantly lower (adjusted *p*<0.05) at baseline vs. TD. However, after week3 (after vancomycin) and week10 (MTT) the relative intensity was observed to significantly increase (adjusted *p*<0.05) and became more similar to TD (Fig. 3A, 5A-B). To quantify the changes, we calculated the AUROC, and sarcosine’s intensity showed 0.82 AUROC at baseline vs TD, at week3 (AUROC=0.73) and at week10 (AUROC=0.85) compared to baseline (Fig. 6A). Notably, the AUROC at week3 and week10 compared to TD were 0.56 and 0.52, respectively (Fig. 6A), which indicates that after MTT, sarcosine’s relative intensity became similar to TD. Similar findings were observed for iminodiacetate (Fig. 6B).

**Figure 5.**
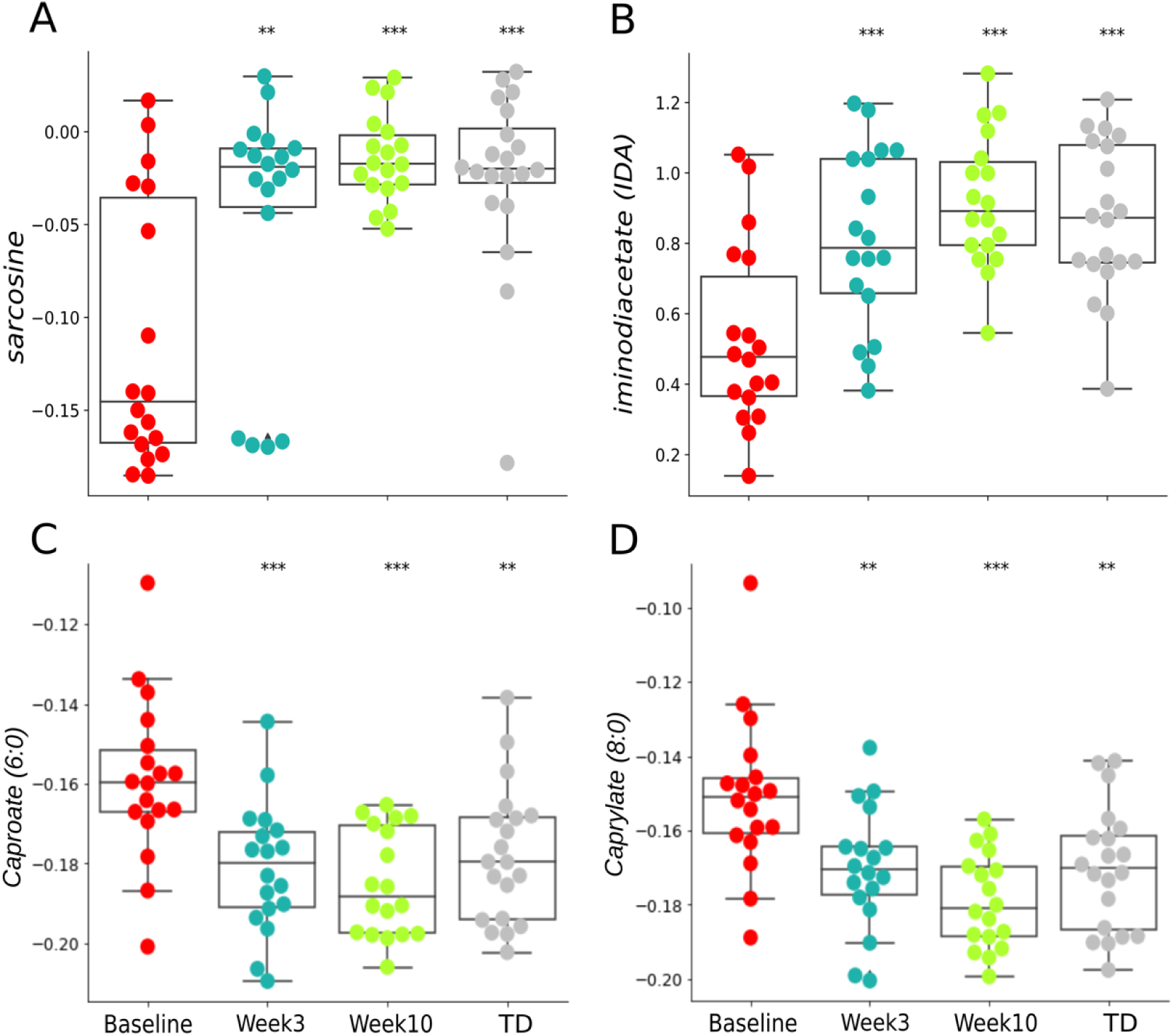
Univariate comparison of the relative intensity (after log10 transformation) in plasma for A) Sarcosine, B) Iminodiacetate (IDA), C) Caproate (6:0), and D) Caprylate (8:0) at ASD baseline vs. MTT (week3, week10) and TD. Each colored dot represents one ASD individual, and grey colored dots represent TD. Asterisks represent significant differences between ASD baseline, and the other groups (*Single asterisk indicates p <0.05, **double asterisks indicate p<0.01, ns not significant, all p-values are FDR corrected). ASD: Autism Spectrum Disorders, TD: Typically Developing.

**Figure 6.**
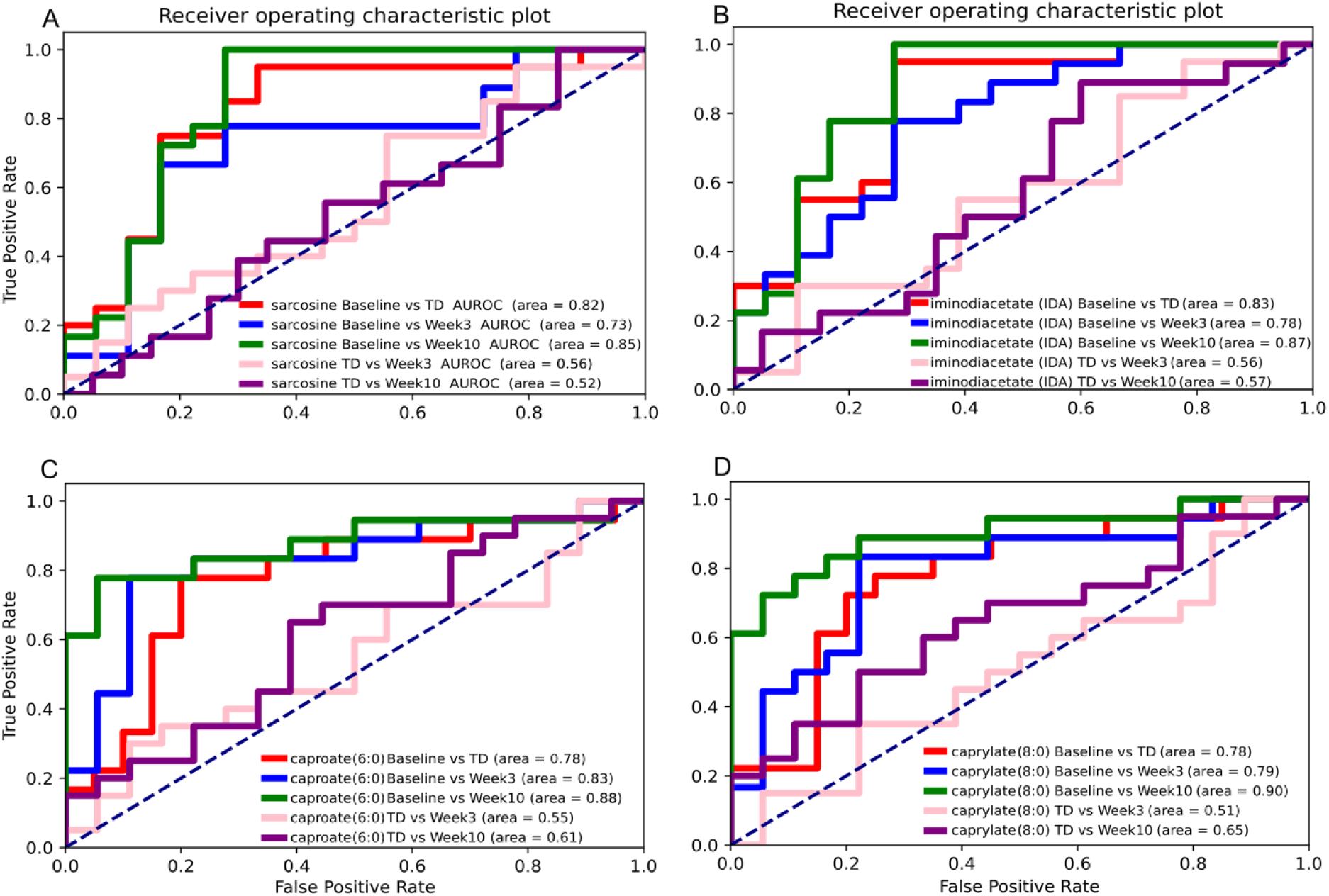
Receiver operating characteristic curves (AUROC) of A) Sarcosine, B) Iminodiacetate (IDA), Caproate (6:0), and D) Caprylate (8:0) for ASD baseline vs. MTT (week3, week10) and TD. Tru positive vs false positive rate was determined across all threshold values to characterize between ASD Baseline against all possible alternative groups.

In contrast, we observed two medium chain fatty acids in plasma, caproate (6:0) and caprylate (8:0), that were significantly higher at baseline vs TD, but after week3 and week10 (MTT) significantly decreased (adjusted *p*<0.05) and their abundances became similar to TD (Fig. 5C-D). Caproate and caprylate AUROCs were 0.78 for baseline vs TD, which showed further differences comparing baseline to week3 (AUROC caproate=0.83, caprylate=0.79) and week10 (AUROC caproate=0.88, caprylate=0.90) vs. baseline, but the AUROC at week3 and week10 vs. TD was between 0.51-0.65 (Fig. 6C-D), which also indicates that after MTT, the relative intensity of caproate and caprylate became similar to TD.

In feces, phenolic compound *p*-cresol sulfate, theobromine, and three sphingolipids; N-palmitoyl-sphingosine (d18:1/16:0), N-stearoyl-sphingosine (d18:1/18:0), and N-oleoyl-sphingosine (d18:1/18:1) were significantly higher at baseline vs TD but after week3 (right after an initial dose of microbiota), week10 (MTT) and week18 (after 8 weeks of MTT) significantly decreased vs. baseline and became similar to TD (Fig 3B, Fig. 7A-D). AUROC calculation showed that baseline vs. TD had 0.76 for *p*-cresol sulfate, after baseline vs. week3 AUROC=0.80, vs week10 AUROC=0.66, vs week18 AUROC=0.75, which indicates after MTT, *p*-cresol sulfate profile changed in children with ASD (Fig. 8A). Interestingly, *p*-cresol sulfate AUROC for TD vs. week3; TD vs. week10; TD vs. week18 were in the range 0.55-0.60 (Fig. 8A), indicating that after MTT, *p*-cresol sulfate peak relative intensity became similar to TD. Similarly, theobromine, N-palmitoyl-sphingosine (d18:1/16:0), N-stearoyl-sphingosine (d18:1/18:0), and N-oleoyl-sphingosine (d18:1/18:1) had higher AUROC after MTT vs. ASD baseline and lower AUROC values after MTT vs. TD, indicating important changes from baseline and greater similarity to TD measurements (Fig. 8B-D). Overall, AUROC validate the findings from univariate analysis and showed the differences between the groups for plasma and fecal metabolites.

**Figure 7.**
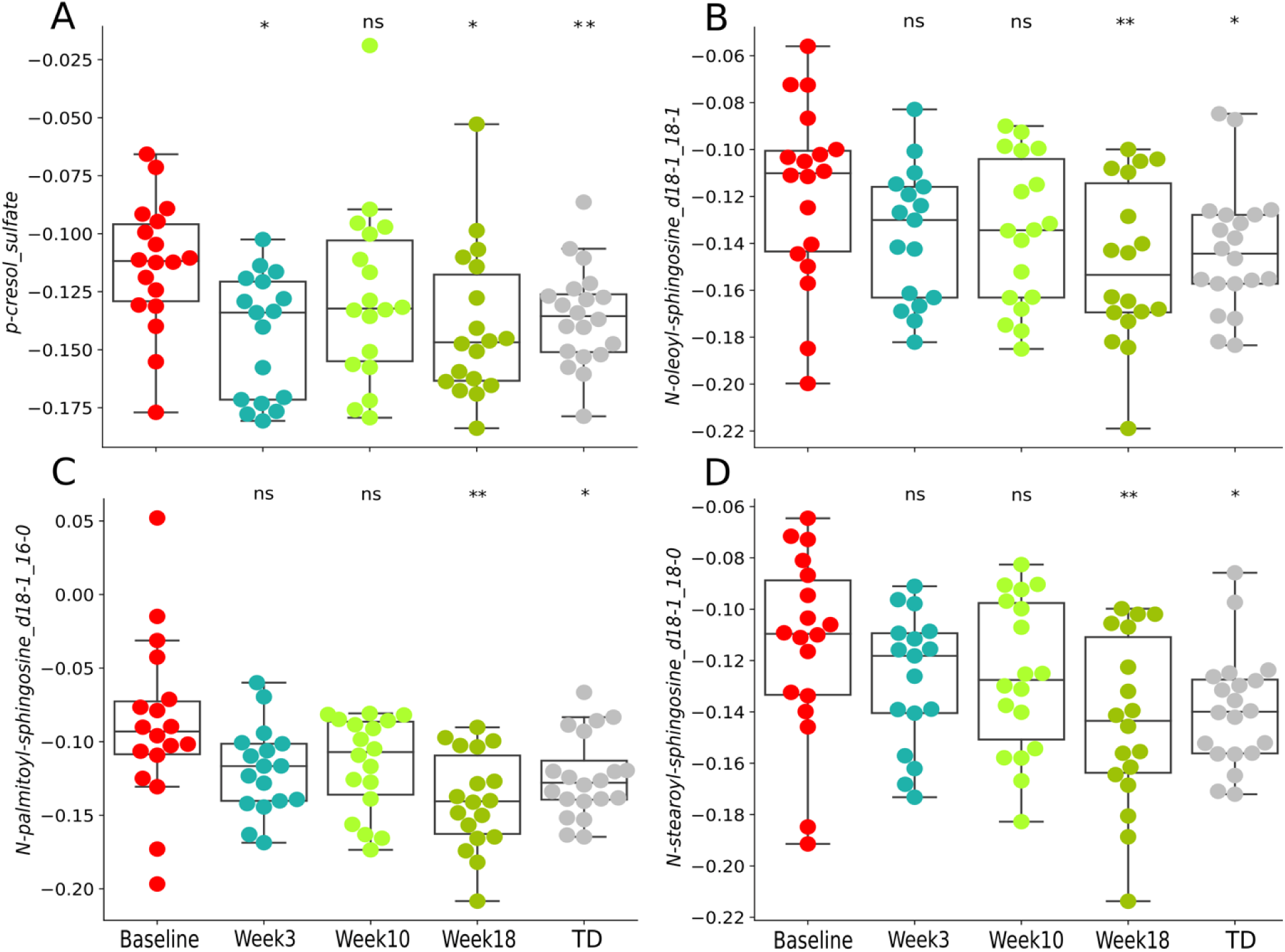
Univariate comparison of the relative intensity (after log10 transformation) in feces for A) p-cresol sulfate, B) N-oleoyl-sphingosine (d18:1/18:1), C) N-palmitoyl-sphingosine (d18:1/16:0), and D) N-stearoyl-sphingosine (d18:1/18:0) at ASD baseline vs. MTT (week3, week10, week18) and TD. Each colored dot represents one ASD individual and grey colored dots represent TD. Asterisks represent significant differences between ASD baseline and the other groups (*Single asterisk indicates p <0.05, **double asterisks indicate p<0.01, ns not significant, all p-values are FDR corrected). ASD: Autism Spectrum Disorders, TD: Typically Developing.

**Figure 8.**
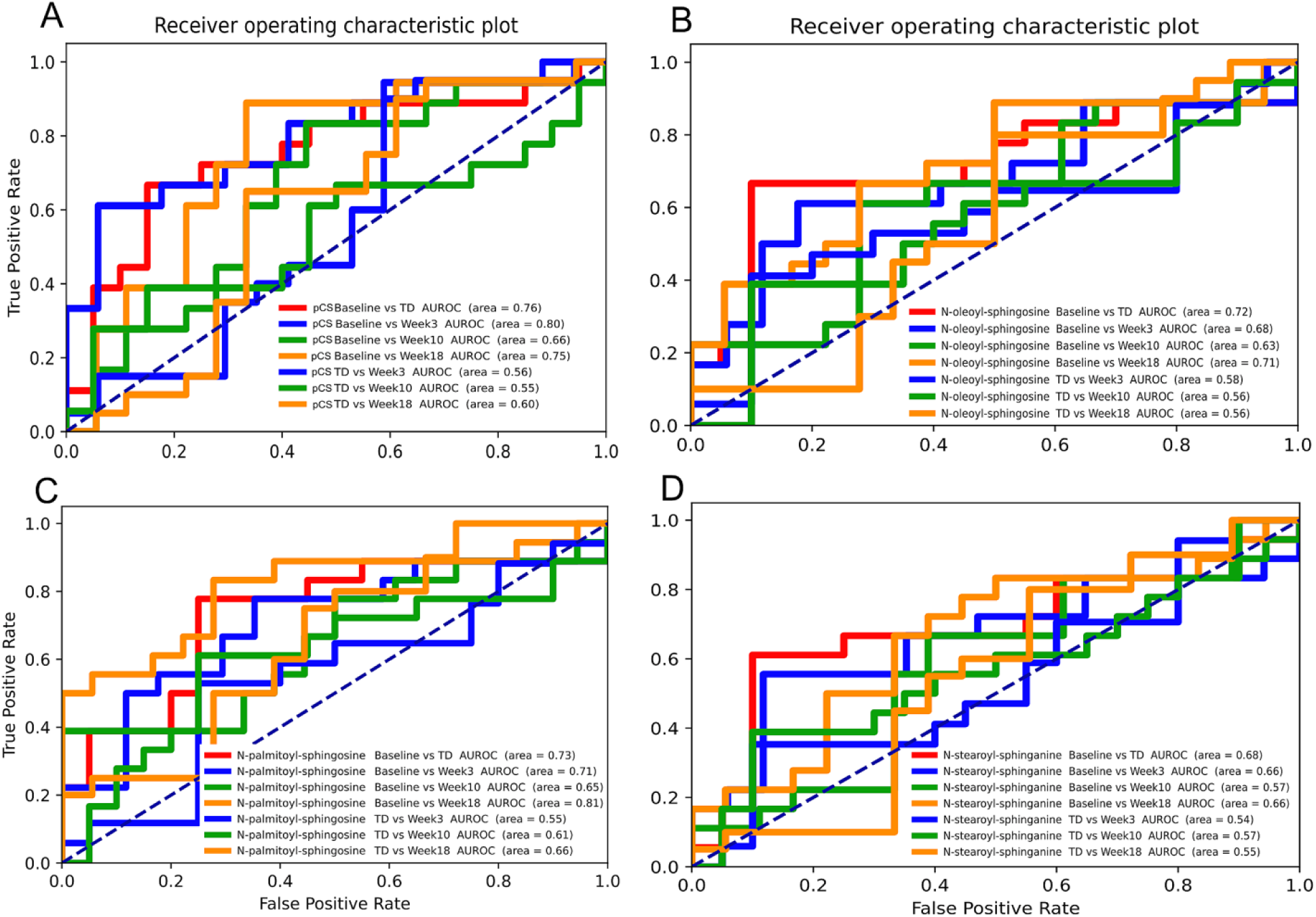
Receiver operating characteristic curves (AUROC) of A) p-cresol sulfate, B) (N-oleoyl-sphingosine d18:1/18:1), C) N-palmitoyl-sphingosine (d18:1/16:0), and D) N-stearoyl-sphingosine (d18:1/18:0) at ASD baseline vs. MTT (week3, week10, week18) and TD. True positive vs false positive rate was determined across all threshold values to characterize between ASD baseline against all possible alternative groups. pCS - p-cresol sulfate.

Recent critiques have questioned whether metabolomic differences between ASD cohorts and controls reflect true tractable differences or arise as a statistical artifact of multiple hypothesis testing, even with traditional test correction approaches (Mitchell et al., 2026). To address this directly, we constructed a null distribution by permuting case-control labels 10,000 times in both plasma and fecal samples, yielding effect size distributions under the null hypothesis of no group differences. Observed metabolomic effect sizes across 42 metabolites significantly exceeded null expectations (p < 0.01), with 92% of features showing stronger separation than permutation maxima. This permutation framework demonstrates that our cohort differences are unlikely to emerge by chance, providing statistical support for the relevance of the observed metabolomic differences (Fig. S8).

### FDA models revealed five key metabolites that notably changed after MTT and became similar to TD

Univariate hypothesis testing identified 20 plasma metabolites and 22 fecal metabolites whose intensities significantly differed between TD and ASD cohorts at baseline or changed after microbiota transplant therapy (MTT) (Fig. 3). Both sets of metabolites were used to separately construct multivariate statistical models *via* FDA using all possible combinations of 3, 4, and 5 variables to distinguish between ASD and TD cohorts. The top-performing FDA model was exclusively made up of plasma metabolites including, 2-palmitoyl-GPC (16:0), caprylate (8:0), laurate (12:0), bilirubin, and indolepropionate achieving an AUROC of 0.99 and a leave-one-out cross-validated (LOO-CV) accuracy of 95%. Similarly, the fecal metabolite data was used to develop an FDA model, with significant features selected through the same univariate approach. The optimal fecal model incorporated *p*-cresol sulfate, hydroxyproline, theobromine, N-oleoyl-sphingosine (d18:1/18:1), and 1-methylhistidine, and similarly achieved a LOO-CV accuracy of 95%.

Utilizing the complete set of 42 significantly differing metabolites from both plasma and feces, all possible 5-metabolite FDA models were also constructed. Through LOO-CV, a model consisting of *p*-cresol sulfate (feces), hydroxyproline (feces), caprylate (plasma), indolepropionate (plasma), and bilirubin (plasma) was identified as the most robust, with a cross-validation accuracy of 95% (Fig. 4D). This model exhibited a type I error rate of 3% and a type II error rate of 0%. Notably, the performance of all multivariate models for classifying between ASD and TD status significantly declined after MTT, as did the measurements taken at the two-year follow-up. The metabolite peak intensities in the model showed significant convergence towards the TD group, with the difference between the median of the five metabolites relative intensity values decreasing by 88% between week0 and week18, compared to the TD group.

### Fecal p-cresol and p-cresol sulfate are positively correlated

We conducted correlation analyses to examine the relationship between certain fecal and plasma metabolites. Fecal *p*-cresol and fecal *p*-cresol sulfate were significantly positively correlated (p<0.001, Pearson-R=0.75) (Fig. S9). Fecal *p*-cresol and fecal *p*-cresol sulfate did not show a significant correlation with plasma *p*-cresol sulfate (Fig. S10-11), which suggests that gut permeability or other factors may affect how much *p*-cresol leaves the gut. *p*-cresol was not detected in plasma, which is not surprising since it has very low solubility in aqueous solutions. We performed correlation analyses with previously published taxonomic and metabolic pathways (KEGG orthologs) (Nirmalkar et al., 2022) but did not observe statistically significant correlations.

## Discussion

A distinct metabolic signature has been consistently observed in plasma and fecal samples collected from children with ASD (Kang et al., 2020; Needham et al., 2021). While several underlying biochemical pathways and environmental factors contribute to these differences, MTT not only improved GI and behavioral symptoms associated with ASD (Kang et al., 2017, 2019) but also leads to a convergence of ASD metabolite profiles towards those observed in TD samples (Kang et al., 2020). Reanalysis of absorbance peaks by Metabolon Inc. using a more comprehensive database (see raw data for plasma and feces in supplemental material SM1), and improvement in our data analysis pipeline allowed for an increase in the number of metabolites detected in plasma and feces (Fig. S3) compared to our previous measurements (Kang et al., 2020). Through reanalysis, we identified 844 metabolites in plasma, and 825 metabolites in feces (Fig. 2), which is a substantial increase in identified metabolites from the original study. Furthermore, several biochemical compounds showed notable shifts in plasma and fecal metabolic profiles after MTT in ASD children. Following treatment, 20 metabolites in plasma and 22 in feces whose peak intensities were initially dissimilar to TD became similar to TD (Fig. 3).

For plasma metabolites, we found similar results to Kang et al., 2020 for bilirubin (an oxidant), iminodiacetate (xenobiotic and a glycine derivative) (Lin et al. 2019), and sarcosine (a glycine derivative, and beneficial against depression and schizophrenia) (Tsai et al., 2004). The relative intensities of these compounds were significantly lower at baseline but increased significantly after MTT in children with ASD (Fig. 2). Also, the intensity of caprylate (8:0, a medium-chain fatty acid) was higher at baseline and significantly decreased after MTT. The analysis for these metabolites shows the consistency of the data previously reported by Kang et al. (2019) and improvement of plasma metabolite profiles after MTT in children with ASD. However, with this re-analysis we detected some potentially important metabolites, such as caproate (6:0, a medium-chain fatty acid), phospholipids such as glycerophosphocholine (GPC), phosphoethanolamine (PEA), glycerophosphoethanolamine (GPEA), and 2-palmitoyl-GPC (16:0). The relative intensity of these metabolites was higher at baseline and significantly decreased after MTT in children with ASD (Fig. 3A). Kang et al. postulated that a decrease in medium-chain fatty acids, such as caprylate to be correlated with an improvement in ASD symptoms (Kang et al., 2020).

Phospholipids are lipids that are highly abundant in neuronal membranes (Joensuu et al., 2020), and they are an important source of energy and provide long-chain polyunsaturated fatty acids (LC-PUFAs). Some research suggests that dysregulated phospholipid metabolism may be a biological component of neurodevelopmental disorders like ASD (Pastural et al., 2009; Tamji et al., 2010; El-Ansary et al. 2011; Brown et al., 2011; Ventura et al., 2020; Needham et al., 2021). Increased sphingolipid-1-PO_4_, a type of phospholipid that was observed in ASD serum compared to TD (Wang et al., 2016). In our fecal samples, relative intensities of sphingosines N-palmitoyl-sphingosine (d18:1/16:0), N-stearoyl-sphingosine (d18:1/18:0), and N-oleoyl-sphingosine (d18:1/18:0) (also called ceramides) were significantly higher at ASD baseline compared to TD but after MTT, they significantly decreased, and their abundance became similar to TD (Fig. 3B). These lipids are important for phospholipid metabolism and mitochondrial function (Tamji & Crawford, 2011). However, the differential abundance of these mitochondrial markers in fecal and plasma metabolites in children with ASD suggests potential mitochondrial dysfunction in children with ASD and may contribute to ASD severity (Needham et al., 2021; Pastural et al, 2009; Rossignol & Frye, 2012; Griffiths & Levy, 2017; Lombard, 1998).

In plasma, GPC (phospholipid) levels were significantly higher in children with ASD compared to their TD counterparts. While GPC levels decreased following MTT, the change was not statistically significant (Fig 3A). Similarly, Needham et al. reported that GPC was significantly higher in ASD children’s plasma samples and was associated with mitochondrial dysfunction (Needham et al., 2021). GPC is a phospholipid-containing choline found in human milk (Dangat & Johsi, 2023) and is an important nutrient for memory, mood, and muscle regulation. However, administration of GPC has been shown to improve cognition against dementia and Alzheimer’s disease (AD) (Amenta et al., 2014; Sagaro et al., 2023). In animal studies, administration of GPC was found to release acetylcholine in rat brains, elevate dopamine and serotonin levels in the frontal cortex, and enhance cognitive function (Traini et al., 2020). Despite the beneficial roles of phospholipids containing choline, increased GPC levels in our samples from children with ASD suggest dysregulation in phospholipid metabolism, and it has been reported that elevated levels of GPC and phosphocholines (PC) can be associated with bipolar disorder patients (Cao et al., 2017).

Another plasma metabolite, indolepropionate (IPA or Indole-3-propionate), was significantly lower at baseline compared to TD (Fig. 3A). IPA is a gut microbiota-derived metabolite of tryptophan, and is increasingly recognized for its neuroprotective and anti-inflammatory properties. In this study, IPA was significantly lower in children with ASD at baseline compared to TD controls, suggesting a potential deficiency in beneficial microbial activity or a disrupted microbial-host interface. Previous studies have reported similarly lower IPA levels in ASD animal models and in children with intrauterine growth restriction (Wang et al., 2023). Interestingly, IPA levels did not significantly change following MTT, indicating that this metabolite may be less responsive to short-term microbial interventions. The persistent reduction (non-significantly) in IPA highlights the need for further investigation into specific microbial contributors and whether longer-term or targeted interventions could restore its production and potential neuroprotective effects in ASD.

In feces, *p*-cresol and *p*-cresol sulfate have consistently emerged as metabolites of interest due to their elevated presence in the urine and feces of children diagnosed with ASD (Altieri et al., 2011, Chen et al., 2013; Kang et al., 2018; Daneberga et al., 2022; Diémé et al., 2015; Emond et al., 2013; Gevi et al 2016; Gevi et al., 2020, Li et al., 2018; Mussap et al., 2020; Osredkar et al., 2023; Perisco & Napolioni 2013, Piras et al., 2022; Tevzadze et al., 2017; Timperio et al., 2022; Flynn et al., 2025, 2026). The primary source of *p*-cresol in the body is its production by gut bacteria, where it act as an antimicrobial compound (Saito et al., 2019). *p*-cresol is produced *via* tyrosine metabolism and is considered a uremic toxin (Candleliere et al., 2022). To eliminate *p*-cresol from the body, the host liver carries out sulfation and forms *p*-cresol sulfate which increases its solubility so that it can be more easily excreted from the body (Gryp et al., 2017; Spanogiannopoulos et al., 2016; Clayton et al., 2009).

We observed that peak intensity of fecal *p*-cresol sulfate was significantly higher at ASD baseline compared to TD (Fig. 3B, 7A). After MTT, *p*-cresol sulfate decreased significantly, and its intensity became more similar to TD (Fig. 3B, Fig. 7A). However, we did not observe significant differences or changes between groups for *p*-cresol in feces, *p*-cresol sulfate for plasma, and *p*-cresol was not detected in plasma. Recent mouse studies have shown that when *p*-cresol was administered, mice exhibited social behavior deficits and more repetitive behavior relative to controls (Bermudez-Martin et al., 2021), and subsequent fecal microbiota transplant from healthy mice led to normalization of ASD-like symptoms (Bermudez-Martin et al., 2021). These findings suggest that *p*-cresol and/or *p*-cresol sulfate may contribute to the pathophysiology of autism. Current results led to a hypothesis that MTT helps to reduce the *p-*cresol sulfate levels by remodeling the gut microbial ecology in ASD children and allowing the growth of microbiota from healthy donors. Another sulfated metabolite that has been associated with ASD in other studies, is 4-ethylphenyl sulfate (4EPS) (Hsiao et al., 2013; Needham et al., 2021; Stewart Campbell et al., 2022). While we did not detect 4EPS in fecal samples, wealso did not observe significant differences or changes between groups in plasma samples. Recent clinical trials in children with ASD using AB-2004, an adsorbent, showed decreased *p*-cresol sulfate and 4EPS levels in urine. These changes occurred alongside improvements in ASD-like behaviors, suggesting 4EPS and *p-*cresol sulfate could be correlated with behavioral severity in ASD (Stewart Campbell et al., 2022).

In feces, another interesting microbial metabolite, Urolithin-A was significantly higher at baseline compared to TD (Fig. 3B). Although Urolithin A is a gut microbiota-derived metabolite produced from dietary ellagitannins found in fruits such as pomegranates and berries (García-Villalba et al., 2013; Selma et al., 2014), its elevated levels at baseline in individuals with ASD may reflect a dysbiotic microbiota enriched in urolithin-producing taxa despite the self-restricted diets commonly observed in ASD. After MTT, the introduction of a more balanced microbial community likely reduced these specialized microbes, leading to lower Urolithin-A levels but no statistical change except at week3 (Fig. 3B). We hypothesize that the high Urolithin-A levels in ASD may be a marker of dysbiosis rather than of beneficial microbial function, and further studies are required to confirm the microbial drivers of Urolithin-A production and its biological relevance in ASD.

It is important to note that certain metabolites such as theobromine and cetirizine (Fig. 3B), while statistically significant between groups, reflect exogenous intake (e.g., chocolate, allergy medications) rather than underlying biological shifts. Their appearance underscores the importance of controlling or at least documenting for diet, medication use, and seasonal variation, which should be explored in future studies with larger, placebo-controlled cohorts and dietary monitoring. A recent critique of metabolomics studies in the context of ASD has posited limitations in reproducibility and biological interpretability more broadly (Mitchell et al., 2026). We directly addressed this criticism by evaluating the robustness of our significant findings, confirming they exceed chance expectations through permutation testing and cross-cohort validation (Fig. S8).

Our most robust multivariate FDA model was composed of five metabolites from either plasma or fecal samples; *p*-cresol sulfate (feces), hydroxyproline (feces), caprylate (plasma), indole-propionate (plasma), and bilirubin (plasma), that were significantly different at baseline vs TD and became similar to TD after MTT (Fig. 3A-B). Fecal *p-*cresol sulfate emerged as a recurring component in many of the top-performing multivariate FDA models. Univariate analysis demonstrated strong differences in the composition of *p*-cresol sulfate between the ASD baseline and TD groups, achieving an AUROC of 0.76. These differences were utilized in the linear multivariate models relying on this variable, highlighting *p*-cresol sulfate as a potentially relevant biomarker for ASD in feces. Higher caprylate (plasma), and lowerindole-propionate (plasma), and lower bilirubin (plasma) were also observed in the previous study (Kang et al., 2020) at baseline and after MTT, relative intensity became similar to TD levels. Indole-propionate, a gut metabolite has been found to be associated with ASD in humans (Yu et al., 2020) and animal models (Wang et al., 2023; Jiang et al., 2024). Hydroxyproline, another FDA model-identified metabolite (Fig. 4D), was previously identified at significantly higher relative intensity in the plasma of children with ASD (Chen et al., 2024).

Combining plasma and fecal metabolite datasets resulted in higher cross-validated accuracy than using either dataset alone, potentially as a consequence of complimenting biological insight provided by each source. The FDA models developed for the reanalysis demonstrated an accuracy of 95%, outperforming previous work with fecal metabolites from the same study, which achieved an accuracy of 94.7% (Qureshi et al., 2020). Previous assessments of plasma metabolites have also shown comparable capabilities to distinguish between ASD and TD cohorts using linear classifier techniques (Howsmon et al., 2017; Adams et al., 2019). This prior work also highlighted that after MTT treatment, the predictive accuracy and average AUROC of fecal metabolites decreased substantially (Qureshi et al., 2020). This pattern was observed in the metabolomic panels composed of both plasma and fecal samples in this study as well, suggesting that improvements in several metabolites may have compromised their predictive validity.Although few large-scale studies have simultaneously examined both plasma and fecal metabolomic profiles across a comparably wide range of metabolites, findings from previously published plasma- or fecal-focused analyses nonetheless provide partial external support for our results. By comparing the relative ratios and directionality of key metabolites across these datasets, emerging consistencies highlight variables that may be particularly informative for model construction. For example, plasma levels of hydroxyproline exhibited moderate discriminability in a comprehensive cohort of 708 children (499 with ASD, 209 TD), aligning with their relative importance in our multivariate framework. These cross-study parallels suggest that, while direct validation remains limited, the identified metabolites capture biologically relevant variance that may generalize to broader or alternative applications (Smith et al., 2023).

While the cross-validation results demonstrate the robustness of these findings, the influence of geographic covariates on microbiome composition should be assessed to determine the broader applicability of the observed associations (Falony et al., 2016). Future studies should incorporate external validation with different cohorts with a large dataset and longitudinal sampling to confirm the reproducibility and clinical applicability of these models. No consistent metabolite alterations were observed across plasma and fecal samples. This may reflect differential host-microbiota interactions, underscoring the complexity of cross-compartmental signaling. Furthermore, studies involving larger cohorts with a placebo arm, along with mechanistic investigations, will be important to better elucidate the metabolic dynamics in ASD and ensure the broader clinical relevance of MTT’s impact.

Although the initial shifts in the metabolic profile observed by week3 reflect the effects of vancomycin treatment, our data suggest that these changes alone do not fully explain the sustained alignment with the TD group seen after MTT. Notably, plasma metabolites remained similar to TD profiles at week18, indicating that MTT may help stabilize or maintain these shifts. Also, we observed only modest changes in fecal metabolites by week3 (after a high-dose of microbiota), with clearer transitions toward TD-like profiles occurring after 8 weeks of MTT. These findings are consistent with Sandler et al. (Sandler et al., 2000), where vancomycin led to temporary improvements in autism symptoms, but these improvements were lost post-vancomycin-treatment, potentially due to lack of recolonization by beneficial microbes. In contrast, our results show that MTT may preserve or reinforce beneficial shifts initiated by antibiotics. This interpretation is consistent with prior findings in recurrent *Clostridioides difficile* infection (Zhou et al., 2023) and highlights the need to consider both antibiotic and transplant contributions in interpreting longitudinal metabolomic responses. In conclusion, the reanalysis of plasma and fecal metabolites revealed an additional 223 plasma and 159 fecal metabolites compared to the previous report by Kang et al., 2020. The contrast in metabolite composition before and after MTT groups provides insight on metabolites with potential biological relevance in children with autism and/or GI issues. This research underscores the value of re-analysis when databases improve and the importance of investigating non-behavioral features in ASD diagnosis and provides valuable insights into the intricate metabolic landscape of this complex disorder.

Using different analysis methods, we consistently observed that sarcosine, iminodiacetate, caproate, and caprylate in plasma were significantly different in ASD vs TD, as reported previously. However, we also identified phospholipids such as glycerophosphoethanolamine (GPE), glycerophosphocholine (GPC), and phosphoethanolamine in plasma, as well as fecal sphingosines N-palmitoyl-sphingosine (d18:1/16:0), N-stearoyl-sphingosine (d18:1/18:0), and N-oleoyl-sphingosine (d18:1/18:0), which were significantly different at baseline compared to typically developing (TD) individuals and/or changed after MTT. Moreover, we observed that fecal *p*-cresol sulfate was higher at baseline compared to TD and decreased significantly after MTT.

Notably, *p*-cresol sulfate in feces was consistently found across our different statistical methods as a potential biomarker before and after MTT in children with ASD. This suggests the potential of microbially-produced metabolite in affecting ASD symptoms. Larger cohort and placebo-controlled trials are necessary to validate our findings and unravel the mechanisms underlying MTT in ameliorating aberrant metabolomic profiles associated with ASD. Furthermore, insights into the underlying biochemical pathways will facilitate the development of more tailored treatment strategies.

An important limitation of this study is the incomplete and partially exogenous-biased coverage of the metabolome provided by the commercial platform, which also contributes to the limited overlap observed between fecal and plasma metabolites. Consequently, the detected metabolite changes represent only a subset of underlying metabolic alterations and should be interpreted within the context of this constrained coverage. While the dataset enables super and subpathway-level and comparative analyses, it may not fully support comprehensive or systems-level biological conclusions. Future studies using broader and more targeted metabolomics approaches are important to extend these findings.

## Supporting information

https://zenodo.org/records/21329932

## Additional Information

The Supplementary File-1 contains Figures S1-S11, raw plasma and fecal peaks data are provided in Supplementary Material-SM1 and difference between old/original and new data are included in Supplementary Material-SM-2-SM5, and they can be found online at:

## Data Availability

The raw LC-MS and LC-MS/MS spectra were generated by Metabolon, Inc. using their proprietary platform. Metabolon does not release raw instrument files or vendor-specific spectra, and therefore these data are not available to the authors. The raw peak-abundance matrix (as provided directly by Metabolon), along with all processed and annotated datasets used in this study, has been deposited in Zenodo (https://zenodo.org/records/21329932). All data necessary to reproduce the analyses in this manuscript are publicly accessible through this repository. These files are also provided as Supplementary materials SM1-SM5.

## Code Availability

Code was developed for analysis purposes for the FDA analysis (Fig.4D), for sample comparison purposes and for permutation testing which can be found at (https://github.com/FatirQureshi/FDA-Model-Search. Zenodo repository associated with this study (https://zenodo.org/records/21329932 and https://zenodo.org/records/21222808) All analyses were performed using standard software packages, as described in the Methods section of the manuscript.

## Abbreviations

4EPS: 4-Ethylphenyl Sulfate
AMP: Adenosine-5’-Monophosphate
ASD: Autism Spectrum Disorder
AUROC: Area Under the Receiver Operating Characteristic Curve
CV: Cross validation
FDA: Fisher’s Discriminant Analysis (context-dependent, could also mean Food and Drug Administration)
FDR: False Discovery Rate
GABA: Gamma-Aminobutyric Acid
GC-MS: Gas Chromatography-Mass Spectrometry
GI: Gastrointestinal
GPC: Glycerophosphocholine
GPEA: Glycerophosphoethanolamine
IMP: Inosine 5’-Monophosphate
KEGG: Kyoto Encyclopedia of Genes and Genomes
LC-PUFA: Long-Chain Polyunsaturated Fatty Acid
LOO: Leave-One-Out
LOO-CV: Leave-One-Out Cross-Validation
MTT: Microbiota Transplant Therapy
PC: Phosphocholine
PCA: Principal Component Analysis
PEA: Phosphoethanolamine
TD: Typically Developing
UHPLC-MS/MS: Ultrahigh-Performance Liquid Chromatography-Tandem Mass Spectroscopy

## Authors Contributions

Conceptualization, K.N., D.W.K., J.B.A., R.K.B.; Data curation, K.N, F.Q., J.B.A., R.K.B.; Formal analysis, K.N, F.Q., J.B.A., R.K.B.; Funding acquisition, D.W.K., J.B.A., R.K.B.; Investigation, K.N, F.Q., J.B.A., R.K.B.; Methodology, K.N., J.H., J.B.A., R.K.B.; Project administration, R.K.B., J.B.A.; Software, K.N.; F.Q.; Supervision, D.W.K., J.H., J.B.A., R.K.B.; Visualization, K.N., F.Q.; Writing—original draft, K.N., F.Q., J.B.A., R.K.B.; Writing—review & editing, , K.N., F.Q., D.W.K., J.H., J.B.A., R.K.B.

## Acknowledgments

We thank all children with ASD and typically developing children for participating in the study as well as the children’s families for their support on the study. We thank Biodesign Center for Health Through Microbiomes Start-up at Arizona State University for the financial support and Joel Meraz for assisting with the references. This work was supported by Finch Therapeutics (formerly Crestovo) to obtain the metabolomics data set.

