## Supplementary material for "Updated Metabolome Annotation Reveals Plasma and Fecal Metabolic Signatures Modulated by Microbiota Transplant Therapy in Autism Spectrum Disorder": https://zenodo.org/records/21329932

**Supplemental Figures S1-S11**

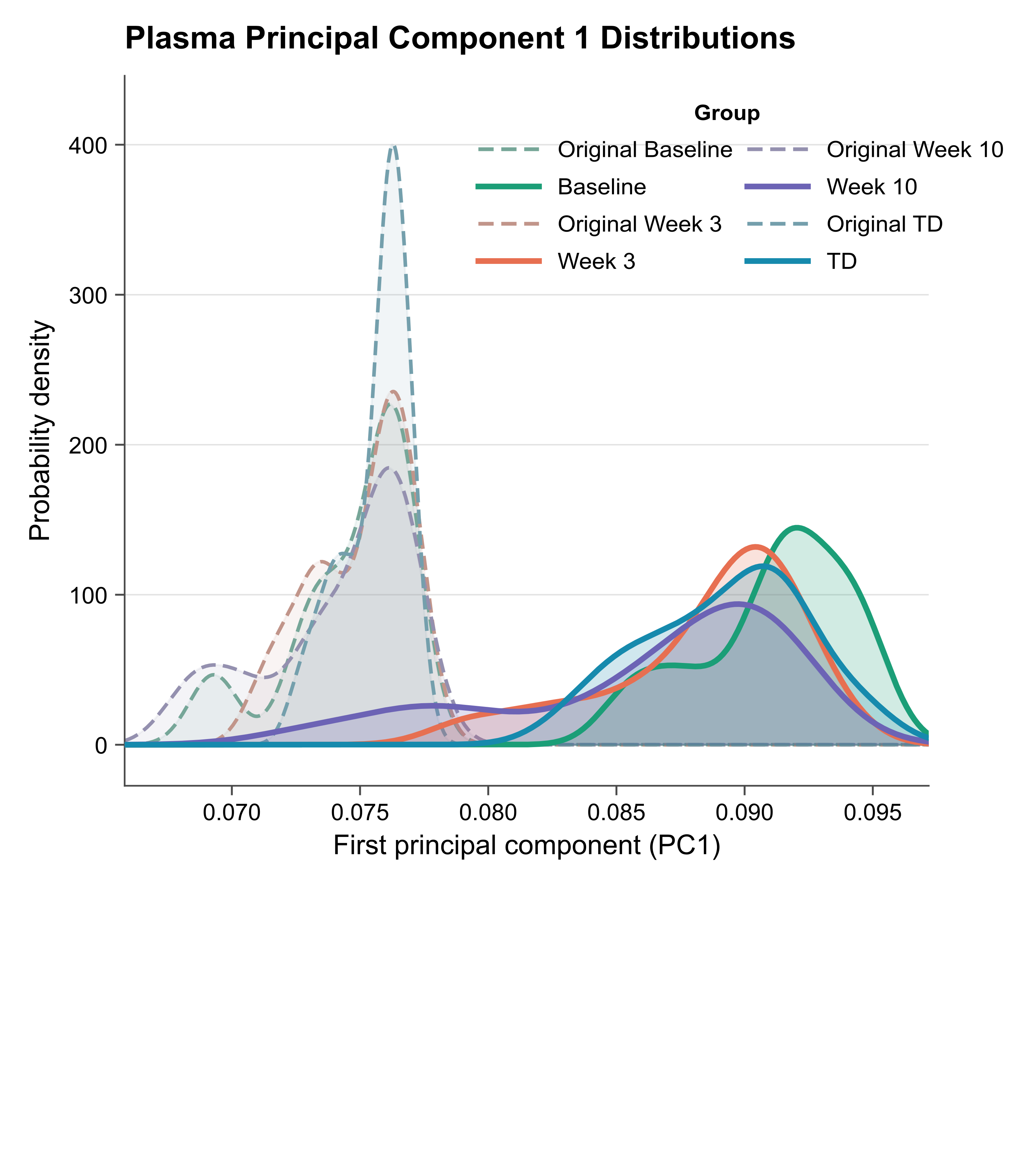

**Figure S1**. **Distribution of plasma metabolite profiles along principal component 1 (PC1) in the original and reanalyzed datasets.** Figure S1. Distribution of plasma metabolite profiles along principal component 1 (PC1) in the original and reanalyzed datasets. Kernel density distributions of principal component 1 (PC1) scores derived from plasma metabolomics data are shown for samples from the original dataset (Original Baseline, Original Week 3, Original Week 10, and Original TD) and the reanalyzed dataset (Baseline, Week 3, Week 10, and TD). PC1 scores were obtained by principal component analysis of plasma metabolite abundance data. The x-axis represents PC1 scores, and the y-axis represents the estimated probability density. Compared with the original dataset, the reanalyzed dataset shows a consistent rightward shift in PC1 score distributions across baseline, treatment, and TD control samples, indicating a systematic change in the global plasma metabolomic profile following data reprocessing and expanded metabolite annotation.

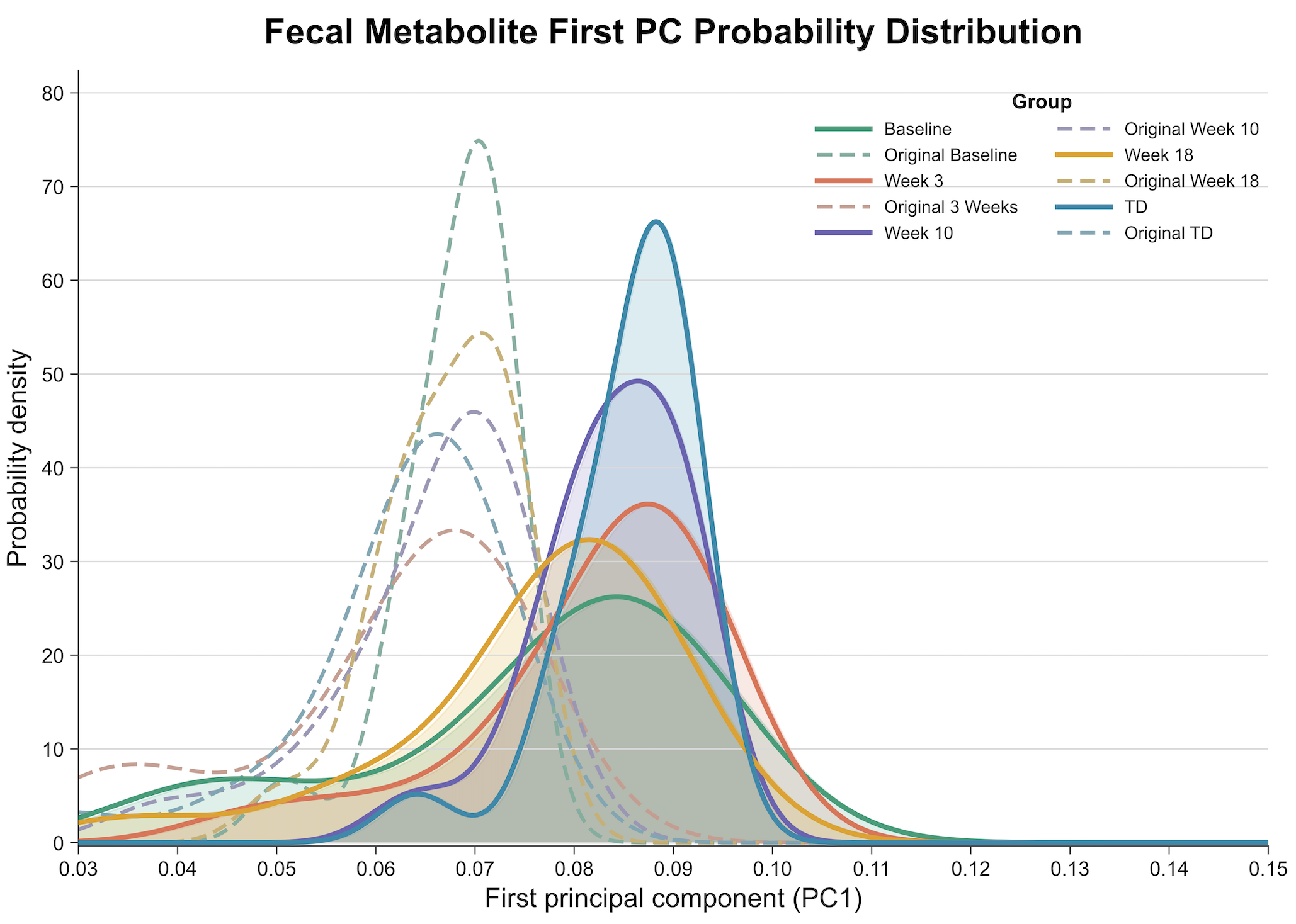

**Figure S2. Distribution of fecal metabolite profiles along principal component 1 (PC1) in the original and reanalyzed datasets.** Kernel density distributions of principal component 1 (PC1) scores derived from fecal metabolomics data are shown for samples from the original dataset (Original Baseline, Original Week 3, Original Week 10, Original Week 18, and Original TD) and the reanalyzed dataset (Baseline, Week 3, Week 10, Week 18, and TD). PC1 scores were obtained by principal component analysis of fecal metabolite abundance data. The x-axis represents PC1 scores, and the y-axis represents the estimated probability density. Compared with the original dataset, the reanalyzed dataset shows a consistent rightward shift in PC1 score distributions across baseline, treatment, follow-up, and TD control samples, indicating a systematic change in the global fecal metabolomic profile following data reprocessing and expanded metabolite annotation.

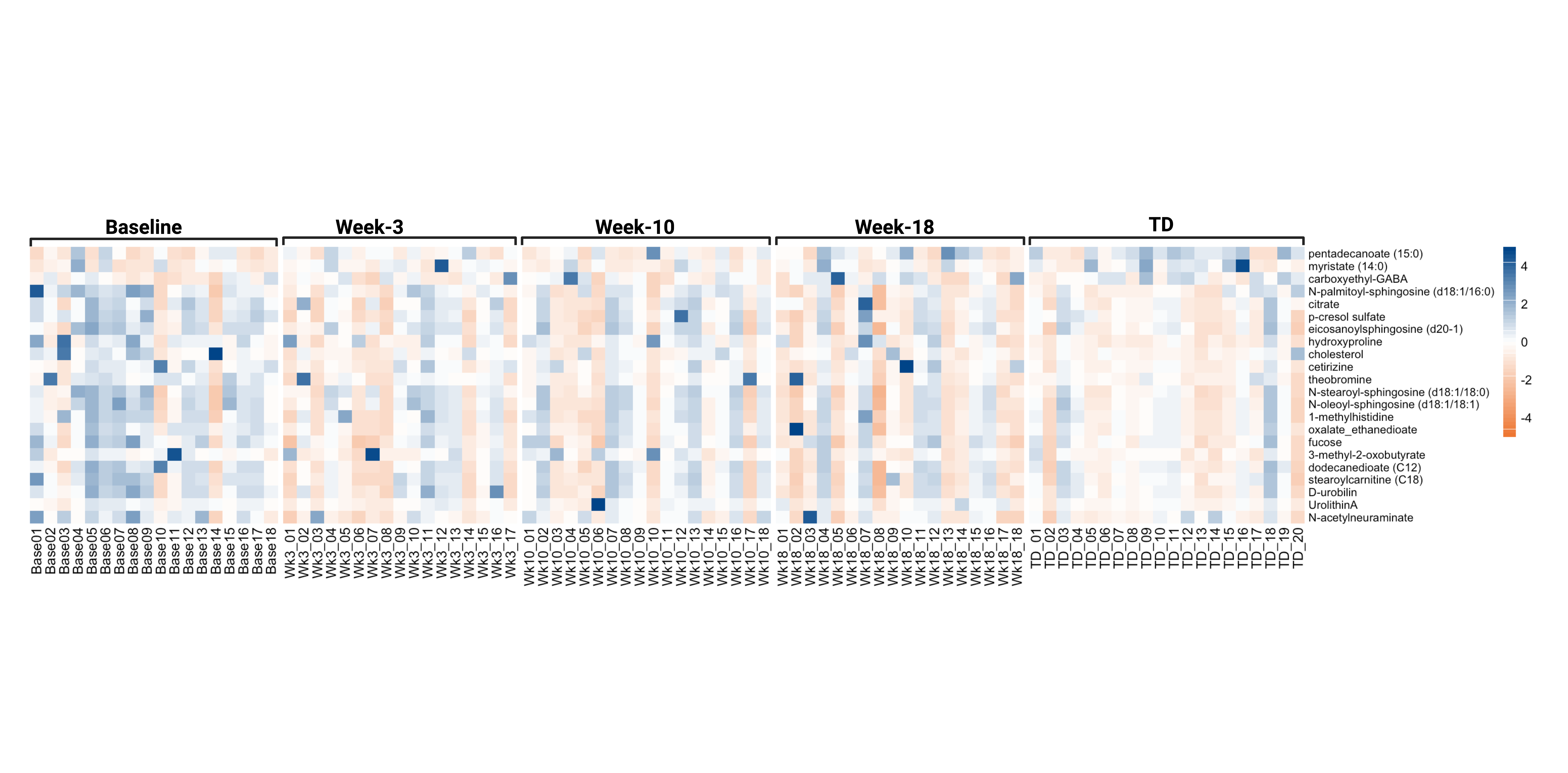

**Figure S3.** Heatmap (z-score) for each participant with 22 fecal metabolites that were significantly changed or were different in children with ASD at Baseline compared with MTT and with TD groups in Figure 3B.

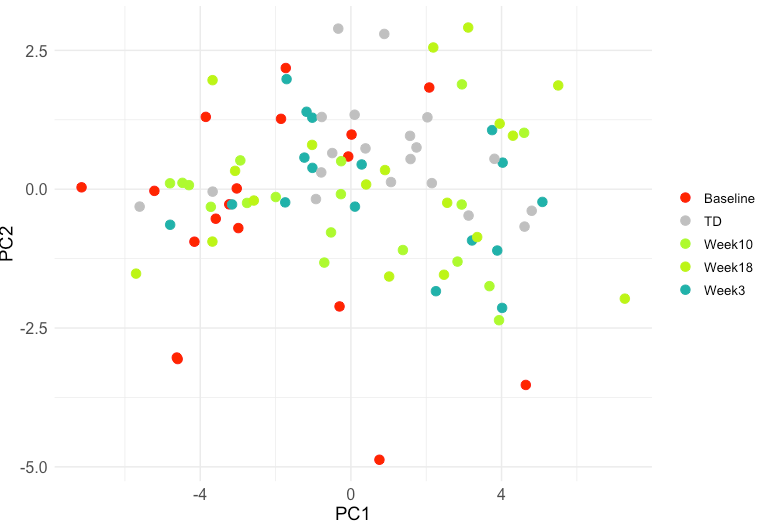

**Figure S4:** Principal component analysis (PCA) of fecal metabolites at Baseline, before and after MTT in ASD, and TD children. ASD – autism spectrum disorder, MTT - microbiota transplant therapy, TD - typically developing.

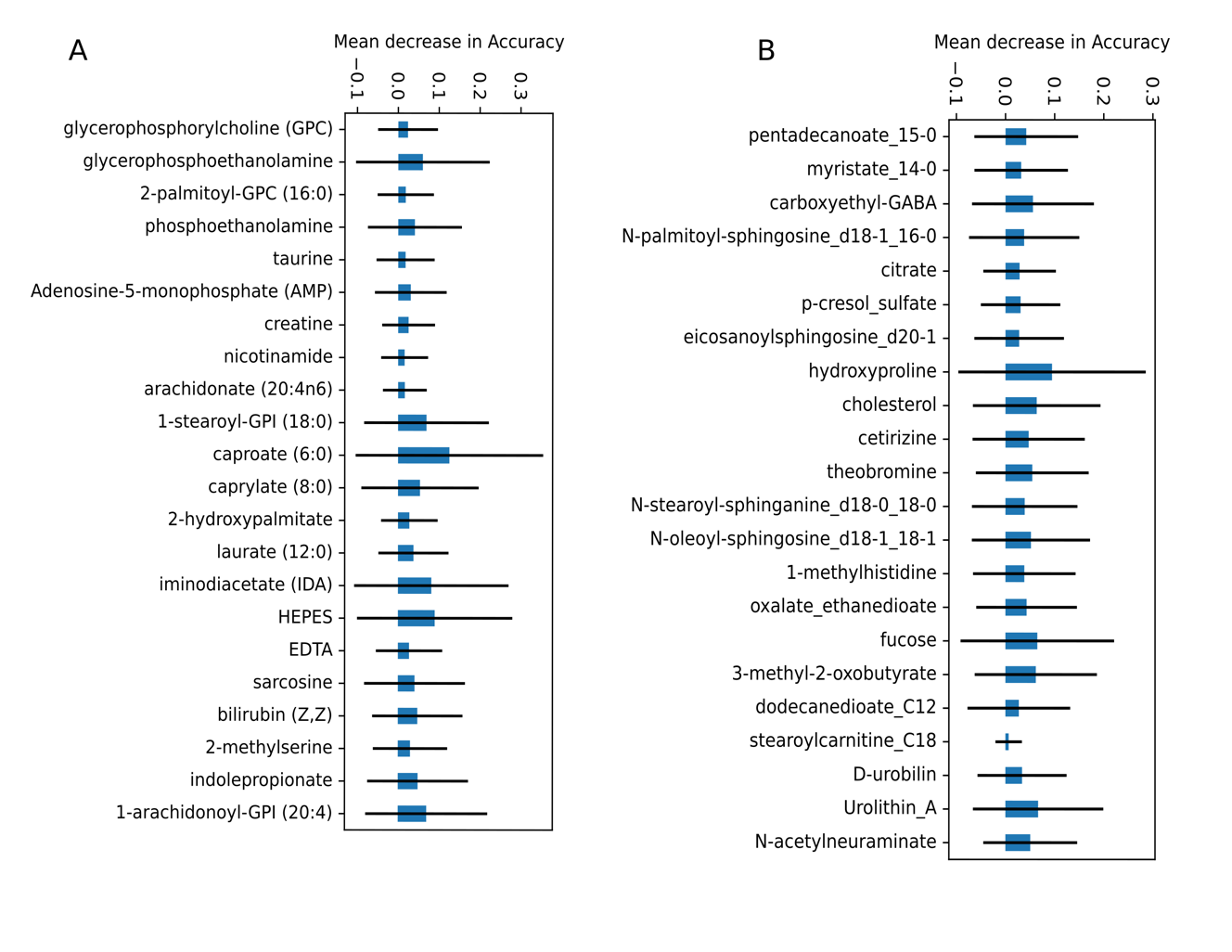

**Figure S5.** The random forest analysis with a mean decrease in accuracy for A) plasma B) fecal metabolites at ASD Baseline vs. MTT (week3, week10, week18) and TD. For the analysis, 22 fecal and plasma metabolites (presented in Figure 2) were used.

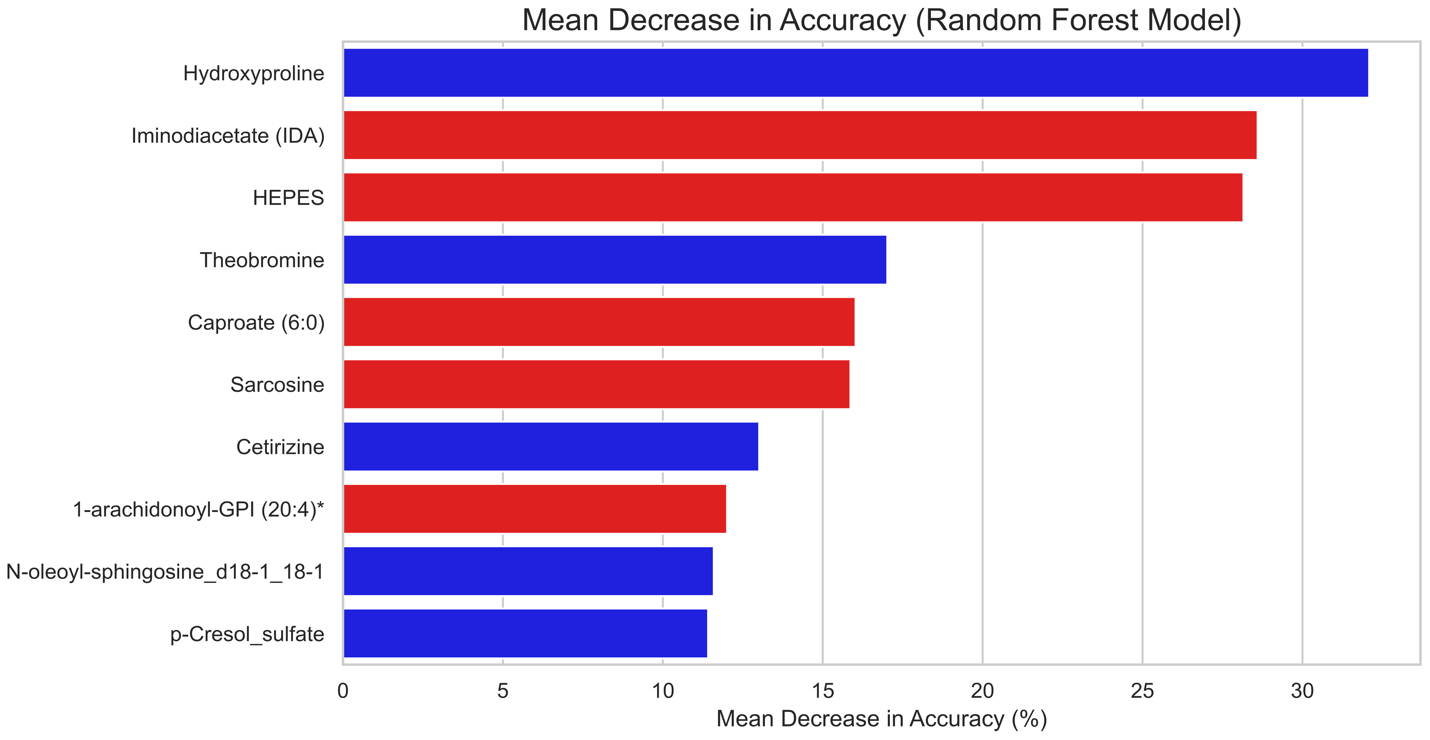

**Figure S6**. Mean-decrease accuracy plot for most significantly changed plasma (red) and fecal (blue) metabolites at ASD Baseline vs. MTT and TD groups. The plots show the random forest analysis, with mean decrease accuracy along the x-axis. Using an ensemble of decision trees, greater mean decrease in accuracy is indicative of a variable playing a greater role in correctly characterizing a sample.

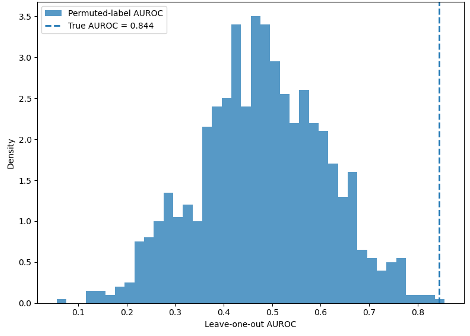

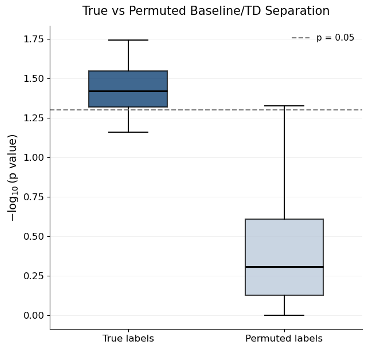

**Figure S8**. Permutation analysis demonstrates robust Baseline/TD metabolomic separation. Baseline and TD labels were randomly permuted 10,000 times while preserving group sizes to generate a null distribution. The observed LOOCV AUROC (left) exceeded most permuted outcomes, and metabolite-level significance values were greater under true labels than under permuted labels (right). These findings indicate that the observed Baseline/TD metabolomic differences are unlikely to arise from random label assignment or nonspecific statistical artifacts.

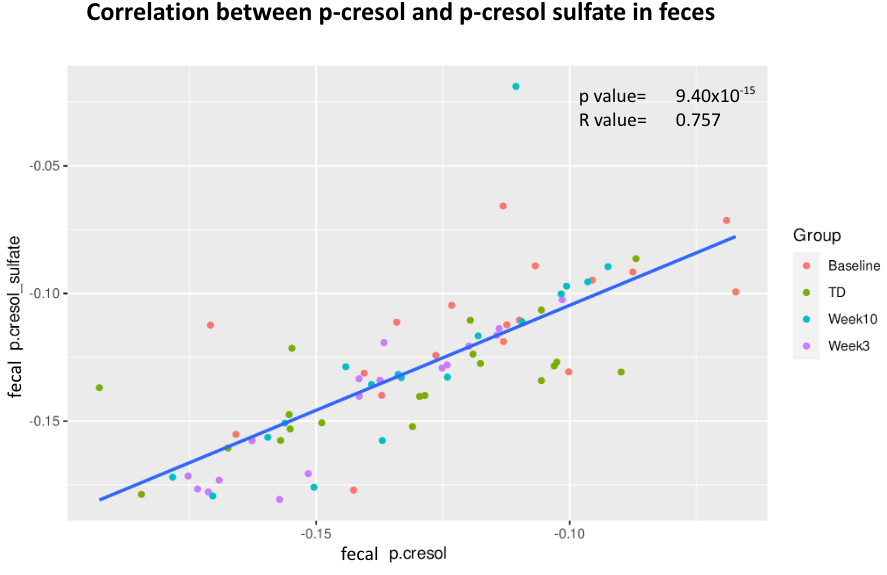

**Figure S9**. Correlation analysis between fecal *p-*cresol sulfate and fecal *p*-cresol between ASD baseline, week3, week10 and TD.

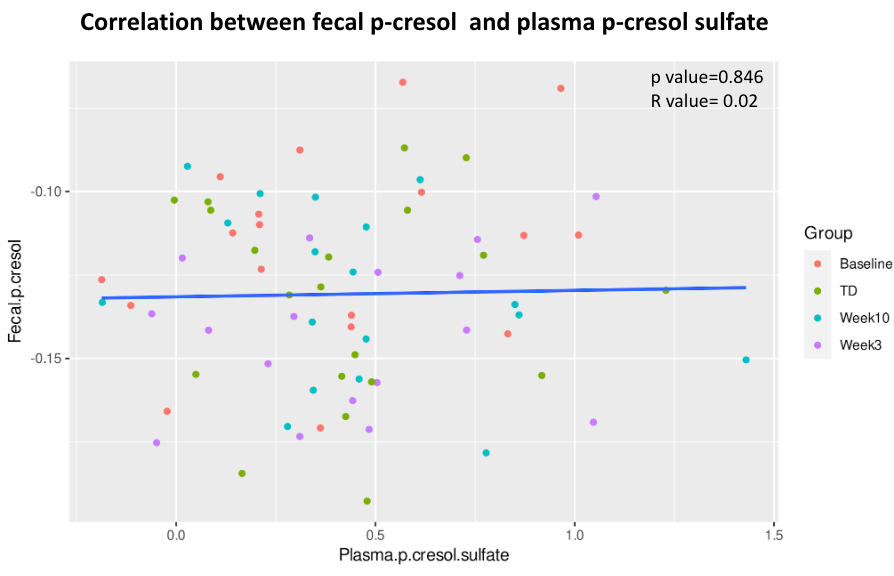

**Figure S10.** Correlation analysis between plasma *p-*cresol sulfate and fecal *p-*cresol sulfate between ASD baseline, week3, week10 and TD.

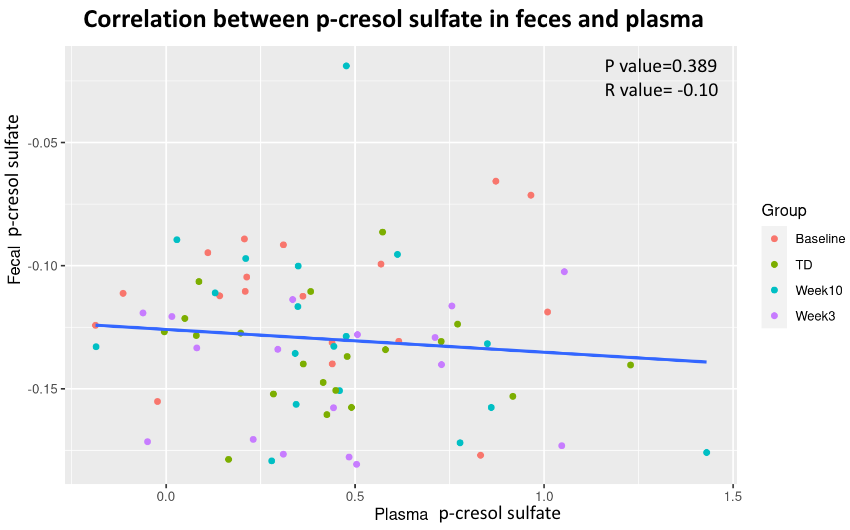

**Figure S11**. Correlation analysis between fecal *p*-cresol and fecal *p*-cresol sulfate between ASD baseline, week3, week10 and TD.
